# Host-specific fungal plant pathogens exhibit distinct interactions with the leaf microbiota of wild grasses

**DOI:** 10.1101/2025.06.24.661283

**Authors:** Victor M. Flores-Nunez, Thaís C. S. Dal’Sasso, Malien Hansen, Graziella Reinhardt, Susanne Braun, Eva H. Stukenbrock

## Abstract

The plant’s microbiome is influenced by the plant species and biotic factors such as the infection by pathogens. Pathogen-microbiome interactions are relevant for the progression of the disease since both can compete within the plant host. We hypothesize that pathogens specialized to different hosts have distinct, direct, and indirect influence on the host microbiome. We focused on the host-specific leaf pathogens *Zymoseptoria tritici* and *Zymoseptoria passerinii*. By using microbiome metabarcoding and coculture interactions, we evaluated the influence of virulent (wild host-infecting pathogen) and avirulent (domesticated host-infecting pathogen) *Zymoseptoria* lineages on the leaf microbiome of the wild grasses *Aegilops cylindrica* and *Hordeum murinum* which are hosts to virulent lineages of *Z. tritici* and *Z. passerinii*, respectively. Our microbiome analysis showed that the fungal communities were affected by virulent lineages, while the avirulent lineages had the most negative correlations with bacteria. Both virulent and avirulent pathogens had the same spectrum of interactions when experimentally cocultured with bacteria. The intensity of pathogen-induced growth enhancement differed between *Zymoseptoria* lineages. We demonstrated that sugar metabolism through the fungal secretion of invertase can be a determinant of bacterial growth enhancement. Our study highlights the role of microbial interactions on host-specificity and mechanisms underlying microbial interactions by *Zymoseptoria* spp.

## Introduction

Plants harbor a diverse microbiome in their tissues that is comprised of bacteria, archaea, fungi, protists, and viruses. These microorganisms form complex associations with the plant, encompassing beneficial, neutral, and pathogenic interactions (1). The plant microbiome plays an important role in facilitating nutrients to the host and reducing the effect of biotic and abiotic stresses (1,2). The factors influencing the plant’s microbiome composition include the plant species and genotype, developmental stage, environmental conditions, the soiĺs microbiome and physicochemical properties, and biological interaction occurring during the infection with pathogens (2,3). Specifically, fungal plant pathogens influence the microbiome diversity and composition by directly interacting with the microbiome members or, indirectly, by influencing the immune response and metabolism of the plant (4).

Fungal plant pathogens specialize to their hosts, this process involves the differentiation of the pathogen’s physical, molecular, and biochemical factors that allow it to infect a particular plant (5). For example, the rapid evolution of genes encoding secreted protein effectors that interfere with the host immune response is a determinant for host specificity in several fungi (6). Moreover, host specialization also involves the mechanisms of tissue invasion, scavenging and uptake of nutrients, reproduction, and coexistence or competition with the microbiota of their specific host (6).

The plant microbiome plays an important role in defence against fungal pathogens by mechanisms that limit their growth and reduce disease severity (1). Pioneer work with the fungal wilt pathogen *Verticillium dahliae* shows that this fungus can manipulate the plant microbiome by secreting antimicrobial effectors to compete against specific antagonistic bacteria during root colonization (7,8) or protect its niche against specific fungi in leaves (9). While some antimicrobial effectors have been co-opted from ancient proteins that evolved before the emergence of land plants (10), others could have a dual function by also interacting with host immune receptors (7). This could imply a role of these effectors in both host-pathogen and pathogen-microbiome co-evolution. Certainly, pathogens evolve, specialize, and diversify together with the associated microbiome of their host, however, the contribution of the microbiome to these evolutionary processes remains poorly understood. Therefore, we ask to which extent pathogen-microbiome interactions can contribute to the evolution of host specificity.

To address this question, we have focused on wild plant-pathogen systems. In this regard, domesticated plant species were shown to have different microbiomes compared to their wild relatives or wild ancestors. Very likely, this reflects the genetic changes in domesticated species following artificial selection in agricultural environments (11). These patterns have been demonstrated in wheat (*Triticum aestivum*), which microbial diversity has been compared to its wild ancestors (e.g., *Triticum dicoccum, Aegilops tauschii,* (12–14)), and in other domesticated grasses like barley (*Hordeum vulgare*, (15,16)). Wild species tend to have more diverse microbial communities in the leaves (17,18) and roots (19), and they also showed a higher extent of selection in the assembly of microbiomes (17). It has been suggested that domestication could have led to the loss of some microorganisms providing important functions for plant growth and defense (20,21). Thus, wild plants could be a relevant source of microorganisms to increase immunity against plant pathogens. Furthermore, pathogens and microbes associated with wild plants represent an intriguing model system to test hypotheses related to coevolutionary dynamics of host-associated microbes and pathogens.

The genus *Zymoseptoria* comprises different grass pathogens including the wheat pathogen *Z. tritici* and has proven a powerful model system for studies of pathogen evolution (22). These fungi are specialized hemibiotrophic pathogens that infect the leaves of wild and domesticated grass species (22). Their specialization has been demonstrated by quantitatively assessing their virulence in different grass species and genotypes under controlled conditions (5,23,24). The pathogens *Z. tritici* and *Z. passerinii* infect the host leaves through the stomata and grow asymptomatically in the apoplastic space (biotrophic phase) for seven to ten days, followed by the induction of plant cell death and the production of asexual fruiting bodies, pycnidia, in the substomatal cavity (necrotrophic phase, (23,25)). Moreover, previous research of the wheat-infecting *Z. tritici* pathogen shows that the infection causes changes in bacterial composition of leaves which is further influenced by the extent of quantitative resistance or susceptibility of the wheat genotype (26). In this study, we focused on two species of *Zymoseptoria* to investigate the extent of specificity of pathogen-microbiota interactions. We specifically asked if microbial interactions may reflect coadaptation and coexistence of leaf-associated microbes and the fungal pathogen in a common host niche.

We have recently described two new *Zymoseptoria* isolates, *Z. tritici* Zt469 and *Z. passerinii* Zpa796, isolated from the wild grasses *Aegilops cylindrica* and *Hordeum murinum* subs. *glaucum*, respectively (5,23). These isolates represent pathogen populations occurring on wild grasses in Iran, and which are close relatives of *Z. tritici* and *Z. passerinii* lineages occurring on the domesticated crops *T. aestivum* and *H. vulgare* subs. *vulgare*, respectively (5,23). The wild host-infecting isolates Zt469 and Zpa796 can produce disease symptoms in the respective wild host (virulent or compatible pathosystem) while domesticated host-infecting isolates, Zt549 and Zpa21, cannot (avirulent or incompatible pathosystem, Supplementary Figure 1A, (5,23)). We hypothesize that due to their specialization to different hosts and microbiomes, virulent and avirulent *Zymoseptoria* lineages will have a distinct, direct and indirect, influence on the host leaf microbiota. Particularly, we expect that the virulent isolates will be more competitive against microorganisms from their own host.

To test these hypotheses, we evaluated the changes in the leaf microbiome induced by the inoculation of virulent and avirulent isolates in a wild host using metabarcoding of the 16S-rRNA and ITS amplicons (Supplementary Figure 1B). We particularly focused on the time when the biotrophic growth of the virulent isolate occurs. To assess which microorganisms were affected by the biotrophic colonization of *Zymoseptoria*, we determined the microbial taxa that correlated negatively and positively with the pathogen. We generated a culture collection to experimentally assess the outcome of pairwise microbial interactions *in vitro*. More precisely, we tested if virulent and avirulent isolates of *Z. tritici* and *Z. passerinii* interact differently with representative members of the wild host’s microbiome by performing *in vitro* confrontation assays (Supplementary Figure 1C).

We identify different impacts on leaf-associated bacteria and fungi during the biotrophic infection of virulent and avirulent pathogens. We further show that virulent and avirulent lineages of *Z. tritici* and *Z. passerinii* do not differ considerably in their interaction with microbiome members of their respective wild host. However, we demonstrate the importance of fungal sugar metabolism on the co-existence of some leaf-associated bacteria which may reflect microbial co-evolution in the plant. Our work sheds light on how the evolution of a specialized plant pathogen in its host has shaped its potential to interact with the plant microbiome.

## Materials and Methods

### *Zymoseptoria* isolates and culture conditions

Two pathosystems were used to determine the influence of virulent and avirulent lineages of *Zymoseptoria* on the leaf microbiome (Supplementary Figure 1A): 1) *Aegilops cylindrica* (Dreschflegel Bio-Saatgut, Witzenhausen, Germany) with the virulent isolate *Z. tritici* Zt469 (isolated from *A. cylindrica*,(5)) and the avirulent isolate *Z. tritici* Zt549 (isolated from *T. aestivum*,(5)), and 2) *H. murinum* ssp. *glaucum* GRA3223 (IPK Genebank, Leibniz Institute, Germany) with the virulent *Z. passerinii* isolate Zpa796 (isolated from *H. murinum*, (23)) and the avirulent *Z. passerinii* isolate Zpa21 (isolated from *Hordeum vulgare* ssp. *vulgare,* (27)). Culture conditions are detailed in the Supplementary Methods.

### Plant propagation and infection assays

*Aegilops cylindrica* and *H. murinum* plants were inoculated with spores of *Zymoseptoria* after 21 and 14 days post-sowing, respectively. Plant infection was performed following the protocol described in (28). Modifications and growth conditions are detailed in the Supplementary Methods.

Samples were harvested by cutting a ∼10cm section of the treated leaf using flame-disinfected scissors. For amplicon sequencing, 5 plants per treatment were sampled, and each leaf was placed in a 15 ml tube with 13 ml of sterile Millipore water. Only mock-treated plants were sampled at 0 dpi (days post-infection), and all the treatments were sampled after 2 dpi during initial plant invasion and the expected biotrophic phase at 4 and 7 dpi. Finally, we also collected samples at the start of the necrotrophic phase at 11 dpi. For microbial isolation, 6 plants per treatment were sampled (at 0, 2, 4, and 7 dpi), and the leaves from the same time point and treatment were pooled in a 15 ml tube with 13 ml of sterile Millipore water. The infection success was evaluated by confocal microscopy (Supplementary Methods).

Samples were processed under axenic conditions. First, the water was decanted from the tubes, and then the leaves were washed with 13 ml of Tween-20 0.02% in PBS 1X (2 min of handshaking) and 13 ml of PBS 1X (2 min of handshaking). Finally, the leaves were pat-dried in sterile absorbent paper tissues before being processed for amplicon sequencing or microbial isolation.

### DNA extraction, library preparation, and amplicon sequencing

The metagenomic DNA from washed leaf samples was extracted with the FastDNA Spin Kit for Soil (MP Biomedicals, Santa Ana, USA). The mechanical lysis and DNA extraction were performed following the procedure described in (26). Library preparation and amplicon sequencing of the 16S-rRNA-V4 and ITS2 markers, for bacterial and fungal microbial communities, respectively, were performed at the University of Minnesota Genome Center (Minneapolis, MN, USA). Library preparation conditions are explained in the Supplementary Methods. Additional sequencing of the 16S-rRNA-V5V7 marker was performed following the protocol reported by Seybold et al., 2020 (26).

### Data processing and analysis

The raw reads of 132 libraries were processed using the pipeline described in Flores-Nunez et al., 2023 (29) with modifications detailed in the Supplementary Methods. After processing, 4,570,240 reads distributed across 116 samples were used for further analysis. All downstream and statistical analyses were performed in R (30), and all plots were constructed using the ggplot2 package (31). The detailed statistical analysis is described in the Supplementary Methods.

### Bacterial and fungal isolation and characterization

To obtain a culture collection of leaf-associated bacteria and fungi that coexist with

*Zymoseptoria* pathogens, we established a culture collection of both bacteria and fungi from *A. cylindrica* and *H. murinum* leaves. Details on the isolation and characterization of isolates are explained in the Supplementary Methods.

### *In vitro* confrontation assays

*In vitro* confrontation assays between pathogen and microbiome isolates were performed to test if virulent and avirulent isolates have different interaction capabilities. All the isolates from the *A. cylindrica*’s culture collection (133) were tested against the *Z. tritici* Zt469 and Zt549 isolates, and from the *H. murinum’*s collection (85) against the *Z. passerinii* Zpa796 and Zpa21 isolates.

The bacterial isolates and four *Zymoseptoria* isolates were confronted in two one-way assays on Fries-3 media 0.5X. For the first assay, a bacterial suspension was mixed with melted Fries-3 0.5X media; after solidification, a suspension of *Zymoseptoria* cells was inoculated on top. In the second assay, a *Zymoseptoria* suspension was mixed with melted Fries-3 0.5X media; after solidification, a suspension of bacterial cells is inoculated on top. A detailed description is given in the Supplementary Methods. Every combination of bacteria and *Zymoseptoria* isolates was tested in an individual Petri dish. Bacteria-agar plates and *Zymoseptoria*-agar plates were incubated at 18°C and 23°C for 7 days, respectively, to give advantage to the microorganism growing on top. Based on the growth of the microorganism in the agar we defined an inhibition phenotype (I) growth enhancement phenotype (P) and no-effect phenotype (N). A detailed explanation of the phenotypes is given in the Supplementary Methods.

We also tested the interaction between fungal species in an *in vitro* assay. The filamentous fungi and isolates of *Zymoseptoria* were used in a two-way confrontation assay. Each fungi-pathogen pair was cultivated together in a single agar plate. The growth of each microorganism in the co-culture was compared with their growth alone in individual agar plates. A detailed description of the design and conditions is provided in the Supplementary Methods. For the yeast-like isolates, we assessed their interaction with *Zymoseptoria* as in the bacterial confrontation assay described above.

The influence of the *Zymoseptoria* lineage on the number of bacteria and fungi exhibiting inhibition, enhancement, and no effect phenotypes was assessed with a Fisher’s exact test.

### Influence of pathogen concentration and sugar on *in vitro* confrontation assays

We determined the range of specificity of the growth enhancement activity of *Z. tritici* on the bacteria isolates isolated from *A. cylindrica* by evaluating the phenotype 1) in the presence of the virulent and avirulent *Z. tritici* and *Z. passerinii* isolates, 2) in the presence of glucose instead of sucrose as the main sugar available in the media and 3) with different cell optical densities (OD). Four different *Pseudomonas* phylotypes that showed growth enhancement in the presence of *Z. tritici* were used for this assay (Supplementary Table 2). The confrontation assays were performed as in the previous section with modifications detailed in the Supplementary Methods. Since the growth enhancement of bacteria did not happen in the presence of glucose, we evaluated the sucrose degradation activity (invertase activity) in the confrontation assays with a 2,3,5-triphenyl tetrazolium chloride (TTC) staining adapted from Lyda et al., 2015 (32) (Supplementary Methods).

### Detection of invertase orthologs in *Zymoseptoria* genomes

Orthologous relationships among proteins from *Zymoseptoria* species were inferred using OrthoFinder v2.2.7 (33), implementing DIAMOND v0.9.24.125 (34) for the all-versus-all comparisons step. Genome data details are explained in the Supplementary Methods. The signal peptide prediction was performed using SignalP 6.0 (35).

## Results

### The wild grasses *A. cylindrica* and *H. murinum* assemble different leaf microbiota

We first addressed the overall difference in microbiome composition of the two crop relatives *A. cylindrica* and *H. murinum* using amplicon sequencing of the 16S-rRNA-V4 and ITS2 regions of bacteria and fungi, respectively. We documented a considerable difference in the microbiome composition of the two plant species propagated in the same soil: Uninfected (mock-inoculated) leaves of *A. cylindrica* showed a higher bacterial and fungal OTU richness and a higher bacterial Shannon index than *H. murinum* leaves (Figure 1A, E; Supplementary Figure 2A, D). In this regard, the host species was a significant driver of bacterial and fungal composition explaining 6.5 and 5.1% of the variation (PERMANOVA, p<0.05, Figure 1C, G), while the sampling time explained 11.9 and 13.7% (PERMANOVA, p<0.05), respectively, (Supplementary Table 3, Supplementary Figure 2B, E), reflecting the dynamic of the microbiota as the leaf matures.

**Figure 1.**
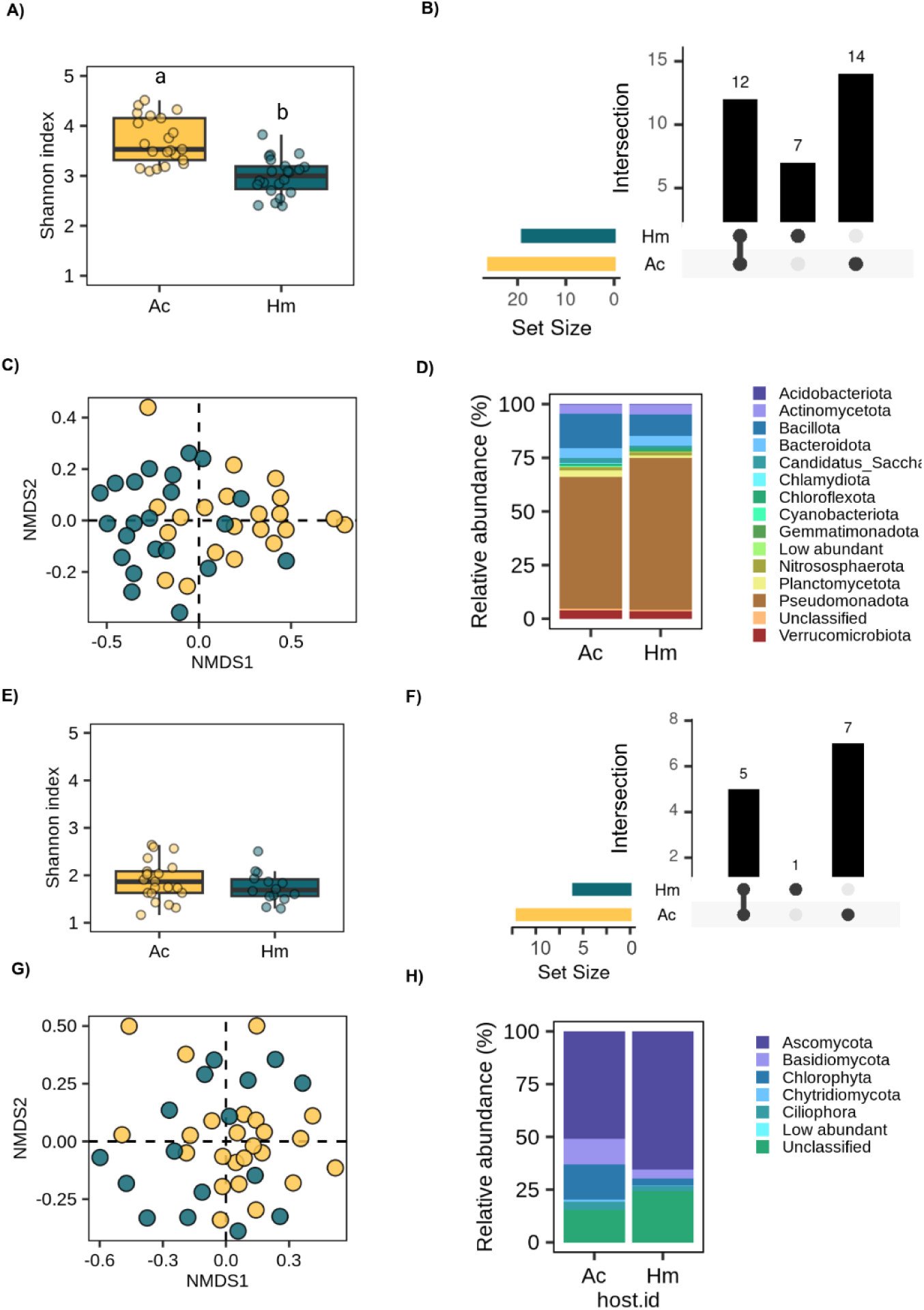
The prokaryotic (A-D) and eukaryotic (E-H) microbiome of uninfected (mock-inoculated) leaves of *Aegilops cylindrica* (Ac) and *Hordeum murinum* (Hm) under greenhouse conditions. The panels show the alpha diversity (Shannon’s index) between hosts (A, E), the intersection of the core OTUs between hosts (B, F), the NMDS based on Bray-Curtis distances of the CSS normalized data between samples (C, G), and the phylum-level classification of OTUs (D, H).

We profiled the taxonomy of the bacterial OTUs as determined by the 16S-rRNA-V4 marker and found that *Pseudomonadota* and *Actinomycetota* were the most abundant prokaryotic phyla in both hosts (Figure 1D). We characterized the bacterial core community of *A. cylindrica* and *H. murinum* mock-inoculated leaves by selecting the OTUs with a relative abundance > 0.1% and a frequency of > 50% in the samples. This allowed us to define a core group of 26 and 19 bacterial OTUs in *A. cylindrica* and *H. murinum* (twelve core OTUs shared, Supplementary Table 4, Supplementary Results) that accounted for 48.7% and 60.6% of the total relative abundance, respectively. Only OTUs from the taxa *Paludisphaera* and *Planctomycetia*, and *Isosphaeraceae* were differentially abundant between the hosts (Supplementary Table 5).

We also profiled the diversity of fungi of mock-inoculated leaves with the ITS2 barcode marker. Hereby, we found that *Ascomycota* was the most abundant group (Figure 1H). We characterized the core fungal communities (relative abundance >0.1, frequency > 20%). Only 12 core OTUs were detected in *A. cylindrica* and 6 in *H. murinum* (five core OTUs shared, Supplementary Table 5, Supplementary Results), which represented a mean of 52.2% and 32.8% of the total relative abundance, respectively. Differentially enriched OTUs between hosts included the basidiomycete yeast *Vishniacozyma* (OTU_17) in *A. cylindrica* and filamentous fungi in the genus Penicillium (OTU_209 and OTU_25) in *H. murinum* (Supplementary table 5).

In summary, we found that *A. cylindrica* and *H. murinum* assembled different leaf-microbial communities when propagated with the same soil-microbial inoculum. However, there was an overlap in dominant core microorganisms between the two plant species. The data from uninfected plants thereby support the hypotheses that *Z. tritici* and *Z. passerinii* encounter distinct microbial species in addition to some common microorganisms during colonization of the host.

### Virulent and avirulent *Zymoseptoria* lineages differently impact bacterial and fungal community composition

We infected plants with virulent and avirulent *Z. tritici* (Zt469 and Zt549, respectively, for *A. cylindrica*) and *Z. passerinii* (Zpa796 and Zpa21, respectively, for *H. murinum*) isolates. We first characterized the infection development using confocal laser scanning microscopy and determined that the virulent pathogens successfully colonized the leaves (Supplementary results, Supplementary Figure 3). The absolute abundance of *Zymoseptoria* reads in the samples reflected that the fungal biomass, as expected, was higher in the virulent interaction compared to the avirulent and mock samples (Supplementary Figure 4).

Based on the PERMANOVA, alpha diversity, and beta-diversity ordination plots (Figure 2A, B, E, F; Supplementary Table 7), we did not find evidence that *Zymoseptoria* confers a strong change in the bacterial community structure in *A. cylindrica* nor in *H. murinum* (PERMANOVA, p>0.05). Conversely, as already observed for the mock-treated plants, the developmental stage of the leaf played a role in the microbiome composition, whereby the sampling time explains approximately 12% and 8% (PERMANOVA, p<0.05) of the variance in *A. cylindrica* and *H. murinum,* respectively. Our secondary sequencing strategy confirmed this observation (Supplementary Figure 5). Consistently, we did not find differentially abundant OTUs between treatments during the biotrophic plant colonization of virulent *Zymoseptoria* or the infection with avirulent isolates.

**Figure 2.**
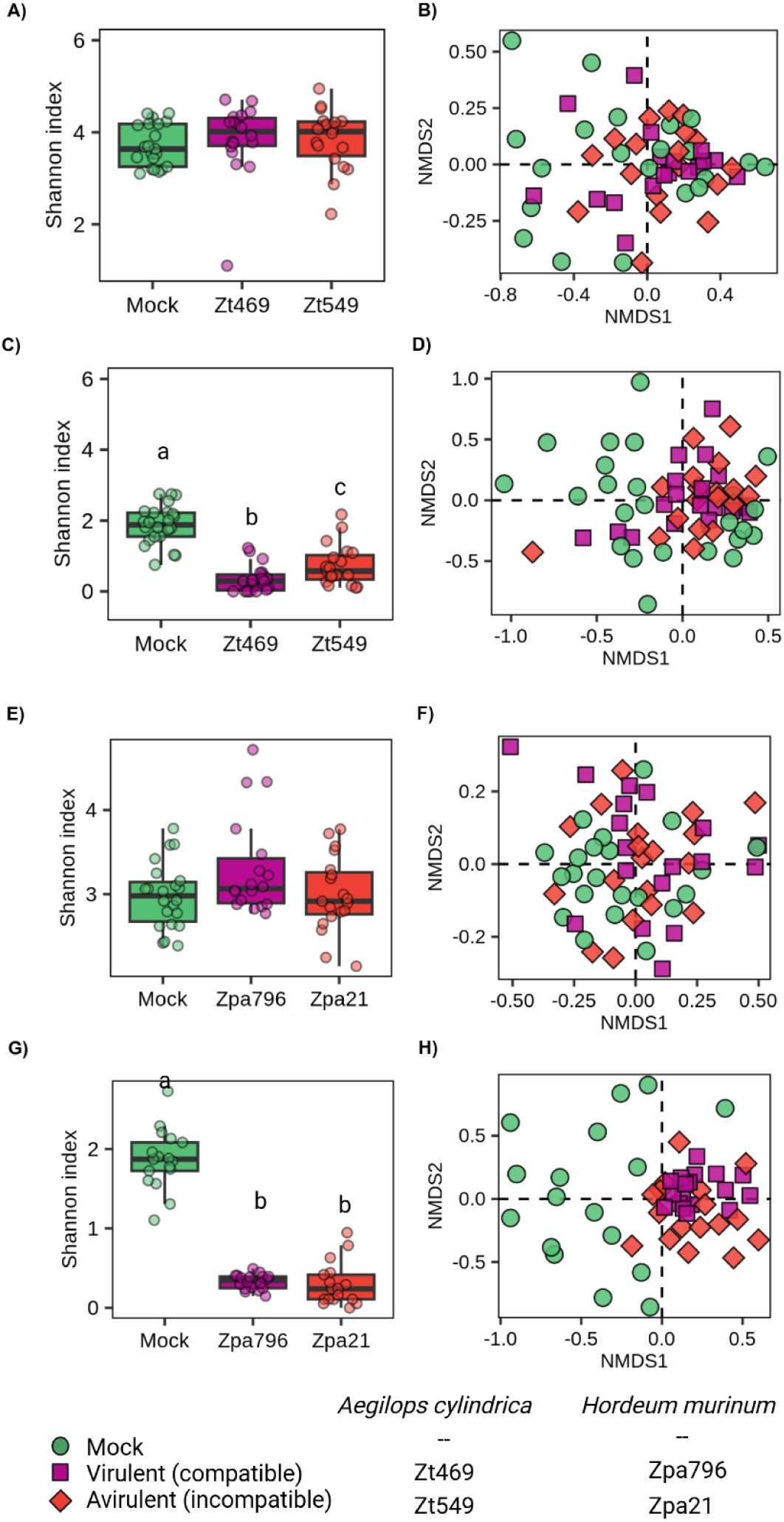
The leaf microbiomes of *A. cylindrica* (A-D) and *H. murinum* (E-H) were infected with different lineages of *Z. tritici* and *Z. passerinii*, respectively. A, B, E, F) 16S-rRNA-V4 barcoding and C, D, G, H) ITS2 barcoding. The boxplots represent the alpha diversity (Shannon index) in the mock, virulent, and avirulent treatments from all the time points while the NMDS were calculated based on Bray-Curtis distances of the CSS normalized data between samples for all time points.

In contrast, the leaves infected with virulent and avirulent *Zymoseptoria* isolates had reduced fungal alpha diversity (Figure 2C, D, G, H) compared to the mock-inoculated leaves (Supplementary Figure 6C, D). Moreover, the infection also influenced the community composition and explained ∼5 and 15% (PERMANOVA, p<0.05) of the variance in *A. cylindrica* and *H. murinum,* respectively (Supplementary Table 7). To differentiate microbial diversity changes caused by pathogen infection and changes due to the dominance of the pathogen reads, we removed *Zymoseptoria* from the OTU table and analyzed the remaining fungal diversity. Doing so, we found that the infection with the virulent isolates, but not the avirulent ones, significantly reduced the fungal alpha diversity. (Supplementary Figure 7). However, a PERMANOVA analysis based on the *Zymoseptoria-*free dataset indicated that the dissimilarity between the samples was not driven by the *Zymoseptoria* treatment (p>0.05).

In summary, the amplicon data of *A. cylindrica* and *H. murinum* indicated that the infection of *Z. tritici* and *Z. passerinii,* respectively, did not affect the leaf-associated bacteria of the two wild grass species. However, the virulent isolates of both pathogens tended to reduce the alpha diversity of leaf-associated fungi.

### Virulent and avirulent lineages of *Zymoseptoria* differentially correlate with members of the leaf microbiome

We next investigated the putative interactions of *Zymoseptoria* with coexisting microorganisms in *A. cylindrica* and *H. murinum*. In order to focus on the most relevant microorganisms, we inferred correlation networks based on the OTU tables for the 4dpi and 7dpi samples, representing the biotrophic host colonization (Figure 3). The correlation network allowed us to identify specific OTUs which are predicted to interact positively or negatively with *Zymoseptoria*. Virulent and avirulent isolates correlate with different core and non-core OTUs (Figure 3A, B, C, D), and 90% of these correlations occurred with bacterial OTUs (Supplementary Table 8, Supplementary Results).

**Figure 3.**
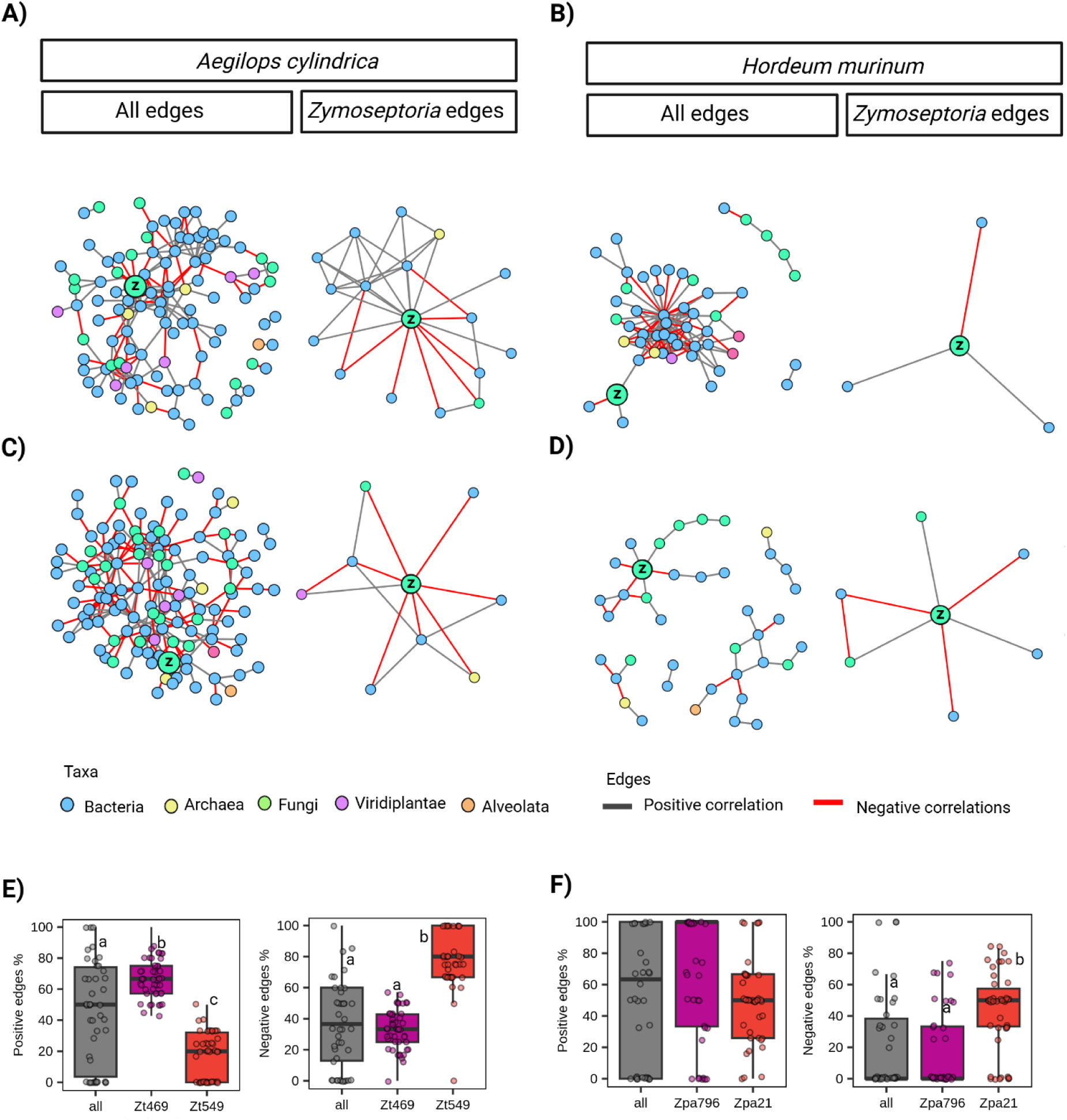
Correlation-network analysis of the *A. cylindrica* (A, C, E) and *H. murinum* (B, D, F) leaf microbiome infected with virulent (A, B) and avirulent (C, D) lineages of *Z. tritici* (Zt469 and Zt549, respectively) and Z. *passerinii* (Zpa796 and Zpa21, respectively) during the biotrophic phase. The “All edges” column corresponds to the complete correlation network in each host and treatment while the “*Zymoseptoria* edges” column corresponds to the subnetwork of the OTUs correlating with *Zymoseptoria*. The boxplots (E, F) represent the proportion of positive and negative correlations of *Zymoseptoria* in 100 randomized networks between the virulent and avirulent treatment, and the randomized joint dataset.

We next asked if virulent isolates can establish more network correlations (positive or negative) with the microbiome members compared to the avirulent isolates. To obtain statistical support for the observed correlations, we computed additional random networks by subsampling the data for each host and treatment, and also for the whole dataset (“all”, Supplementary Methods). Then, we compared the proportion of positive and negative correlations in the whole data set against the proportion of correlations in the specific treatments (virulent and avirulent) and hosts. Positive correlations occurred considerably more frequently in the virulent Zt469 (mean=67.13%) than in the avirulent (mean=16.64%) Zt549 infection in *A. cylindrica* when compared to the joined datasets (mean=45.76 %, Figure 3E). The number of positive correlations found for Zt469 surpassed almost twice the average OTU connectivity in the networks (Supplementary Results, Supplementary Figure 9). Moreover, we observed more negative correlations for the avirulent interactions in both *H. murinum* and *A. cylindrica* (Figure 3E, F). In summary, the different correlation patterns between isolates suggest a different influence on the microbial communities by virulent and avirulent isolates of both *Z. tritici* and *Z. passerinii*.

### Different intensity of pathogen-microbiome interactions in *A. cylindrica* and *H. murinum* leaf microbiome

To further disentangle the relevance of direct and indirect interactions between *Zymoseptoria* pathogens and their host microbiomes, we generated a culture collection of bacteria and fungi from *A. cylindrica* and *H. murinum* grown under controlled conditions in an agricultural soil (see Methods and Supplementary Results). To test if virulent and avirulent *Zymoseptoria* lineages have different capabilities to interact with the host microbiome, pairwise *in vitro* confrontation assays were performed. We designed the confrontation assays to identify antagonistic, neutral, and beneficial interactions among the fungal pathogens and the respective microbiome members of the host (Figure 4), i.e. between *Z. tritici* and the *A. cylindrica* isolates (117 bacteria and 24 fungi), as well as between *Z. passerinii* and the *H. murinum* isolates (59 bacteria and 16 fungi).

**Figure 4.**
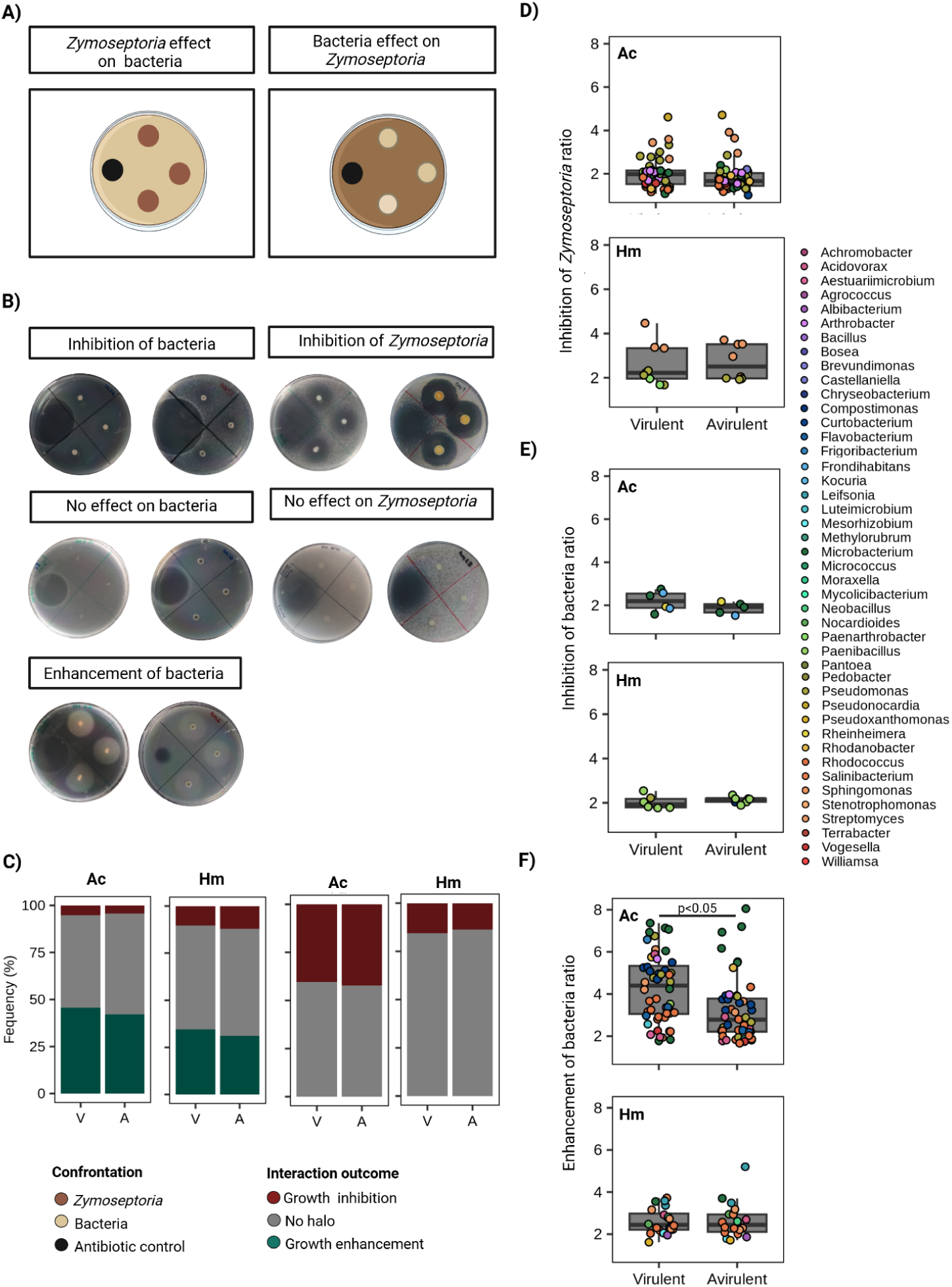
Confrontation assays of the cultivable leaf-associated bacteria of *A. cylindrica* (Ac) and *H. murinum* (Hm), with the virulent (V) and avirulent (A) *Z. tritici* and *Z. passerinii* lineages, respectively. A) Experimental setup. B) Representative pictures of the outcomes in each assay. C) The frequency of each interaction outcome with the virulent and avirulent *Zymoseptoria* lineages with the leaf-associated bacteria. The boxplots represent the intensity of the interaction outcome measured as the ratio between the inhibition or enhancement halo and the top colony. D) Inhibition of *Zymoseptoria* by bacteria, E) Inhibition of bacteria by Zymoseptoria, and F) growth enhancement of bacteria by *Zymoseptoria*.

More than 40% and 25% of the tested bacterial isolates inhibited the growth of Z*. tritici* and *Z. passerinii*, respectively. Particularly, *Pseudomonas* and *Streptomyces* isolates have the strongest inhibiting effect on the fungal pathogens as revealed by the size of the inhibition halos (Supplementary Figure 10 and 11). The rest of the bacterial isolates did not show any effect on the growth of *Zymoseptoria* (neutral interactions), but they were able to grow on the media with the pathogen (Figure 4B, C).

Interestingly, *Z. tritici* and *Z. passerinii* were only able to inhibit a smaller number of bacterial isolates (∼5 and ∼10%, respectively), including *Penibacillus* and diverse *Actinobacteria* (e.g. *Paeniarthrobacter*, *Microbacterium*, etc) (Figure 4B, C; Supplementary Figure 10 and 11). Surprisingly, a diverse array of bacterial genera from four phyla showed enhanced growth in the presence of *Z. tritici* and *Z. passerinii* (∼45 and ∼40%, respectively, Figure 4B, C; Supplementary Figure 10 and 11). This growth enhancement was particularly prominent among isolates of *Pseudomonas*, *Rhodococcus*, *Acidovorax*, *Microbacterium*, and *Curtobacterium*.

We compared the proportion of antagonistic, beneficial and neutral interactions of the virulent and avirulent fungal isolates. Hereby we find no significant differences in the frequency of confrontation phenotypes between virulent and avirulent *Zymoseptoria* isolates (Figure 4C, Supplementary Table 9). This assay indicates that the general outcome of the *in vitro* confrontation between *Zymoseptoria* and bacteria is not different between virulent and avirulent isolates.

Based on the intensity of the interaction phenotype (defined as the size of the inhibition or growth enhancement halo), the *A. cylindrica* infecting isolate Zt469, in general, induces a stronger enhancement compared to the wheat infecting isolate Zt549 (Figure 4D), particularly for bacteria of the genera *Pseudomonas*, *Rhodococcus*, *Curtobacterium,* and *Streptomyces* (Supplementary Figure 12). For the two *Z. passerinii* isolates, there was however no difference in the strength of bacterial growth enhancement or inhibition between the virulent Zpa796 and the avirulent Zpa21 (Figure 4E, F, G). This finding suggests that *Z. tritici* lineages might possess differential activity to enhance the growth of leaf-associated bacteria.

We next performed a set of confrontation assays with the collection of fungal isolates from *A. cylindrica* (26) and *H. murinum* (16). Interestingly, filamentous fungi showed the same interaction outcomes, ranging from growth inhibition and enhancement when co-inoculated with *Zymoseptoria* on a petri dish (Figure 5A, B). However, the most common phenotype was the growth inhibition of *Z. tritici and Z. passerinii* isolates induced by diverse filamentous fungi from the two host plants (∼50-90% and ∼75-90%, respectively). The isolates with the highest inhibitory activity belong to the genus *Penicillium* and the genera *Bjerkandera* and *Terfezia (*only for *Z. passerinii*, Supplementary Figure 13). Both *Z. tritici and Z. passerinii* can inhibit a few filamentous fungi too (∼10% and ∼30% of fungal isolates, respectively), with *Penicillium* isolates (phylotype 1_OTU209) the most consistently inhibited. Only a few filamentous fungi from the *A. cylindrica* culture collection showed enhanced growth when co-cultured with *Z. tritici* (∼5%, Figure 5C), including isolates of the genera *Phialemonium*, *Aspergillus,* and *Apiotrichum* (Supplementary Figure 13). Finally, the virulent Zt469 was significantly inhibited by more fungal isolates than the avirulent Zt549 (p>0.033, Supplementary Table 9).

**Figure 5.**
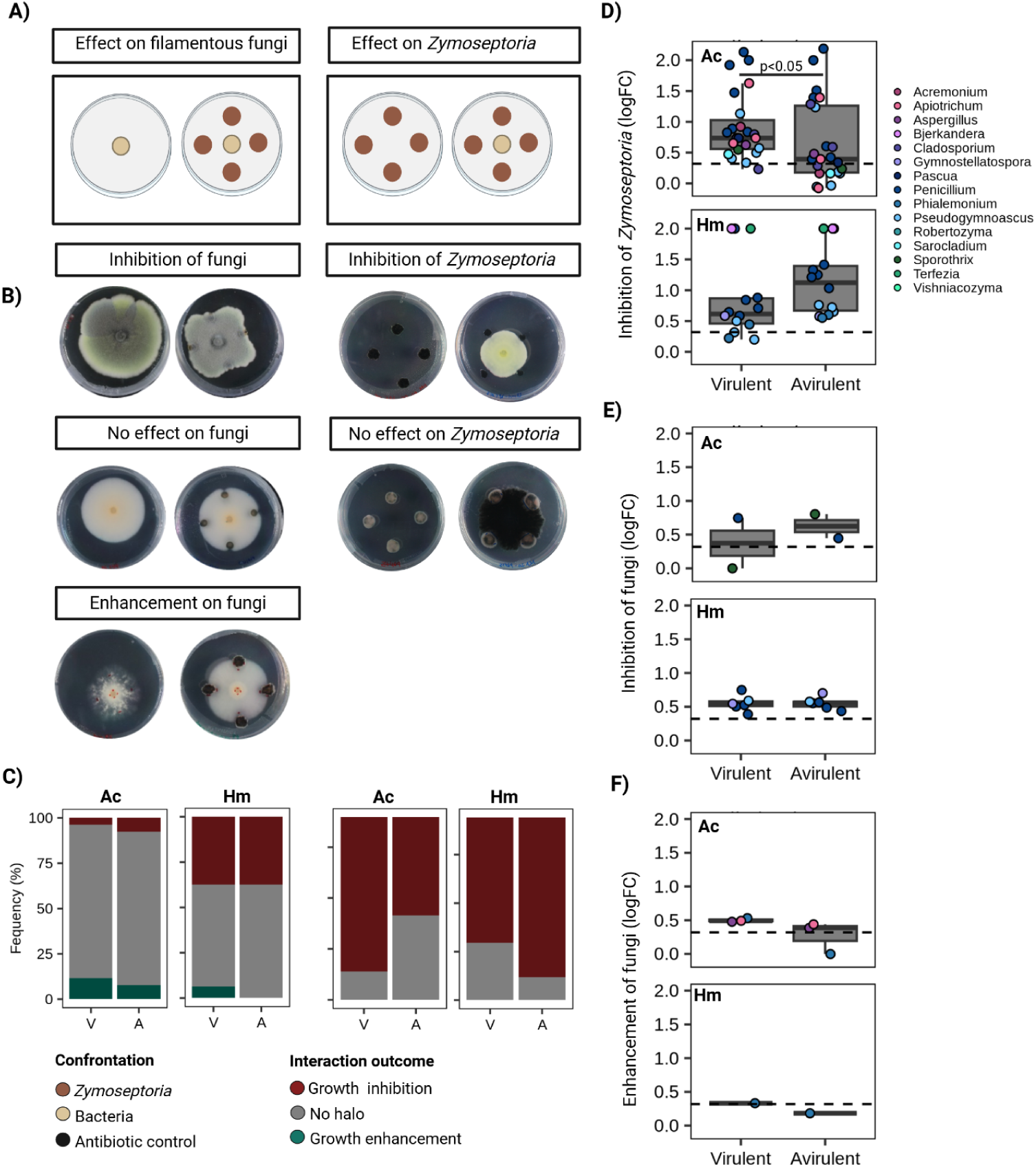
Confrontation assays of the cultivable leaf-associated fungi of *A. cylindrica* (Ac) and *H. murinum* (Hm), with the virulent and avirulent *Z. tritici* and *Z. passerinii* lineages. A) Experimental setup. B) Representative pictures of the outcomes in each assay. C) The frequency of each interaction outcome with the virulent and avirulent *Zymoseptoria* lineages with the leaf-associated fungi. The boxplots represent the intensity of the interaction outcome measured as the logFC between the diameter of the isolate growing in confrontation and a isolate growing alone. D) Inhibition of *Zymoseptoria* by fungi, E) Inhibition of fungi by Zymoseptoria, and F) growth enhancement of fungi by *Zymoseptoria*.

### Pathogen-derived sugar and cell concentration define the growth enhancement phenotype of bacteria

Since the outcome of a culture-dependent confrontation test can be affected by the culture conditions, we conducted a more detailed investigation of the putative conditions underlying the fungus-microbe interactions. To this end we assessed the role of the pathogen species, pathogen load, and main sugar source on the confrontation outcomes. We focused on leaf-associated *Pseudomonas* from *A. cylindrica* as the *Pseudomonas* bacteria showed the highest diversity of phylotypes and confrontation outcomes (Supplementary Figure 10). To evaluate the *Pseudomonas-Zymoseptoria* interactions, we focused on the particular isolates with the growth enhancement phenotype, which was induced in different intensities by the two *Z. tritici* isolates (Figure 4, Supplementary Figure 12, Supplementary Table 2).

The virulent Zt469 consistently enhanced the growth of four *Pseudomonas* phylotypes (ac26, ac30, ac66, and ac9), while the avirulent Zt549 enhanced the growth of only two bacterial isolates. *Z. passerinii* (Zpa796 and Zpa21) did not enhance the growth of any of the *Pseudomonas* isolates, although Zpa796 enhanced *Pseudomonas* ac9 (Figure 6a,b). In order to determine the role of the fungal cell density, we inoculated the pathogen with two different OD values. Hereby, we find that a smaller amount of pathogen inoculum on the media (from 2.7 to 1.35 OD) reduces the size of the growth enhancement halo of the *Pseudomonas* bacteria (Figure 4a). Based on this observation, we conclude that a certain quorum of cells is needed for this growth-enhancement phenotype.

**Figure 6.**
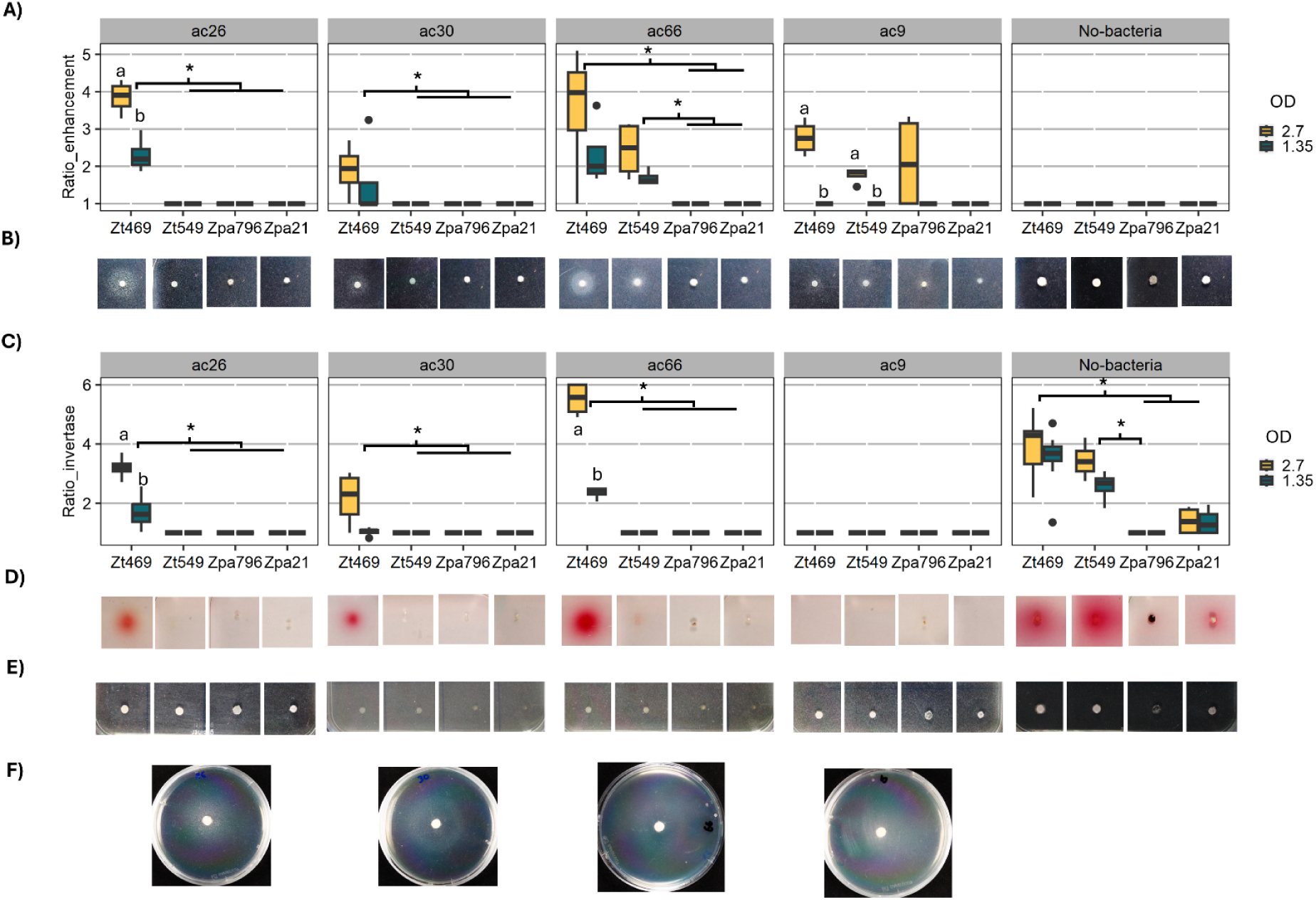
The growth enhancement activity of *Z tritici* and *Z. passernii* with *Pseudomonas* phylotypes isolated from the leaves *of A. cylindrica.* A) Confrontation test of 4 *Pseudomonas* phylotypes and 4 *Zymoseptoria* isolates inoculated at 2 different optical densities (OD) in sucrose-containing agar media, the ratio between the halo size divided by the colony size is represented. B) Representative pictures of the growth enhancement phenotypes with *Zymoseptoria* isolates at 2.7 OD. C) TTC staining of reducing sugars during the confrontation in sucrose-containing media, the ratio between the red halo and the colony size is represented. D) Representative pictures of the TTC staining with *Zymoseptoria* isolates at 2.7 OD. E) Confrontation test of 4 *Pseudomonas* phylotypes and 4 *Zymoseptoria* isolates inoculated at 2 different optical densities (OD) in glucose-containing agar media. F) Growth enhancement phenotype induced by 10 uL of glucose 2M. The lowercase letters indicate a p-value < 0.05 between cell OD (wilcox-test) while the asterisk indicates a p-value < 0.05 between pathogen isolates (Kruskall-Wallis test with Dunn post-hoc test).

We further addressed the possible mechanisms whereby *Zymoseptoria* can enhance growth of bacteria in its surrounding. Previous transcriptome analysis of the *Z. tritici* reference isolate IPO323 showed an up-regulation of a secreted invertase enzyme when growing on culture media with sucrose (36). Based on this observation, we hypothesized that our *Zymoseptoria* isolates secrete an invertase that breaks sucrose into monosaccharides providing a nutrient source for co-occurring bacteria which otherwise cannot utilize the sucrose in the media. To test this, we first tested if different carbon sources in the media would impact the bacterial growth enhancement. We tested the growth of bacteria co-inoculated with *Zymoseptoria* in a medium with glucose instead of sucrose. With the glucose plates, we find an absence of the growth enhancement halo otherwise present in the sucrose plates (Figure 4e). To further test the hypothesis, that the available sugar determines the growth enhancement of *Pseudomonas* bacteria, we conducted a further experiment: In order to mimic the release of glucose by *Z. tritici*, we added a paper disc soaked with a glucose solution on the top of an agar plate containing the bacteria. The bacteria had an enhanced growth surrounding the glucose-paper disc, indeed suggesting that the bacteria grows better in the presence of this monosaccharide (Figure 6f). Finally, we determined if extracellular invertases of *Zymoseptoria* were active in the media by staining the resulting monosaccharides with TTC. Interestingly, *Z. tritici* had the biggest diffusion of reducing sugars in the control agar media compared to *Z. passerinii,* as evidenced by the bright red halos (Figure 6c,d). The invertase activity remained active in the virulent *Z. tritici* Zt469 confronted with the bacterial isolates, but not in the avirulent Zt549 or *Z. passerinii*.

We next performed an orthology assessment in our *Zymoseptoria* isolates to determine if the fungi possess orthologous of the IPO323 (the reference strain) secreted invertase (Supplementary table 9). Our analysis showed four different groups containing orthologous beta-fructofuranosidases (EC3.2.1.26). Each isolate encoded at least one potential invertase copy with a predicted secretion signal. We conclude that the pathogen isolates have the potential to secrete invertases in the media or during plant infection. Based on the in vitro assays with *Pseudomonas*, we conclude the bacterial growth enhancement is due to the hydrolysis of sucrose in the media and its expression might depend on the microbial context.

## Discussion

### The infection of *Zymoseptoria* spp. induce differential changes in leaf-associated bacterial and fungal communities

This work is conceived with the concept that *Zymoseptoria*, a genus of host-specialized fungal pathogens, evolves in the context of host specific microbiota. Indeed, plant species and genotype are important factors that influence the plant’s microbiome composition (17,37). In our pathosystems, the wild *A. cylindrica* and *H. murinum* hosts showed a small but significant species differentiation with common and unique core leaf microbiome members (Figure 1). This supports the notion that *Z. tritici* and *Z. passerinii* have coexisted and evolved with specific microorganisms in the leaves of their hosts. The microbiome composition that we have studied under controlled greenhouse conditions might be different from the microbiome in the field or natural environment (38), however we consider that conserved functional traits of these microorganisms still may define a component of the core *A. cylindrica* and *H. murinum* microbiomes.

We have previously described lineages of *Z. tritici* and *Z. passerinii* that have specialized in their wild host (Zt469, Zpa796) and diversified from the lineages that infect domesticated relatives (Zt549, Zpa21(5,23)). Here, we framed these lineages in the context of the wild host, as virulent lineages (for the adapted wild host-infecting isolates) and avirulent lineages (for the non-adapted domesticated host-infecting isolates). We focused on the wild pathosystems because domesticated crops such as bread wheat and barley have been shown to assemble different and less defined microbiomes from their wild relatives (13,15,17). Therefore, it is possible that the pathogens have evolved with different microbiomes in each host (domesticated and wild) leading to different patterns of interactions.

Fungal plant pathogens influence the microbiome of their host plants (4). Our data shows that the leaf-associated bacterial communities of wild grasses are more resilient to the changes induced by virulent and avirulent pathogens than the fungal communities, which showed a reduced alpha diversity when infected with the virulent isolates (Figure 2). These changes occurred in the asymptomatic biotrophic phase of infection. A similar effect was described in *Arabidopsis thaliana* leaves infected with the biotrophic pathogen *Albugo* spp. (39). Moreover, Seybold et al. (26) showed that the modulation of the host’s immunity and metabolism by *Z. tritici* also affected the bacterial leaf microbiome of wheat. Our network analysis suggests that the virulent Zt469 correlates positively with more OTUs (Figure 3), which may reflect an indirect positive effect on bacteria that could be promoted with the induction of systemic susceptibility (ISS, (26)). Similarly, the higher proportion of negative correlation by the avirulent Zt549 and Zpa796 might indicate a decrease in abundance due to pathogen-triggered immunity (PTI,26)) against the pathogens. The transcriptomic survey of Zt469 (virulent) and *Z. tritici* IPO323 (avirulent) in *A. cylindrica* shows differential expression of pathogen resistance genes supporting that defense genes indeed are differentially induced during fungal infection (40). The suppression or induction of immune related genes may indirectly influence the colonization of other microbial taxa in the leaf tissue.

Microbial interactions may also be direct between the invading fungal pathogen and the host microbiota. We used in vitro confrontation assays as a proxy for direct mechanisms of interaction. *Zymoseptoria* species can inhibit and enhance the growth of different bacteria and fungi isolated from the wild host, suggesting their potential to directly influence or modify the host microbiome (Figure 4 and 5). The *in vitro* antifungal activity of *Zymoseptoria* against fungi like *Penicillium* and others does not explain, by itself, the reduction of other fungal taxa. Pathogen-derived antifungal molecules might have specificity for certain groups of taxa, as observed for an antifungal effector in *Verticillium dahliae* (9). The reduction of diversity could also be associated with the dominance of the pathogen on the leaves, which could cause a displacement of the other fungal taxa (41). The production of antimicrobials to remove microbial competitors is a recent field of study that remains to be explored in *Zymoseptoria* species. Currently, a couple of effectors with antimicrobial activity have been described in *Z. tritici* (42,43), which represent potential candidates to study in the context of the host microbiome. In this study, we generated a collection of antagonistic bacteria and fungi from wild wheat and barley which can be further explored for putative wheat biocontrol consortia for control against *Zymoseptoria tritici*.

### Direct pathogen-microbiome interactions show signatures of host specificity

It has been suggested that the ability of pathogens to coexist or compete with other microbes inhabiting plant surfaces and tissues is an important factor for its host specificity (6). We used the in vitro confrontation assays to test if virulent and avirulent specialized *Zymoseptoria* lineages would establish different interactions with the host leaf microbiota. *Zymoseptoria tritici* isolates specialized to wild and domesticated wheat have the same interaction outcome when confronted with the leaf-associated bacteria from the wild host (Figure 4). The same pattern is true of isolates of *Z. passerinii* from wild and domesticated barley. This observation indicates that the pathogens might have a similar repertoire of metabolites or effectors to influence the host microbiome. *Aegilops cylindrica*-infecting Zt469 and *Triticum aestivum*-infecting *Z. tritici* IPO323 show high synteny levels across the core chromosomes (44). Similarly, the *Zymoseptoria* isolates show the same susceptibility against the antifungal repertoire of the host leaf-associated bacteria. Indeed, the biocontrol activity of microorganisms against pathogens is broad, and a single species might have the potential to antagonize diverse fungal pathogenic species (45). However, we found that more fungal isolates inhibit the virulent Zt469 than the avirulent Zt549, which indicates a possible difference in susceptibility towards other host-associated fungi. Previous studies have indeed demonstrated that the biocontrol fungus *Trichoderma harzianum* can inhibit different pathogen species, and have different expression patterns with different co-occurring pathogens (46).

Differences in the expression of effectors have been described in isolates *Z. tritici* with different infection patterns and virulence in wheat (25), for example in the antimicrobial ribonuclease Zt6 (25,42). Interestingly, similar ribonucleases produced by *Ustilago maydis* (Ribo1) and *Cholletotrichum fruticola* (CfRibo1 and CfRibo2) are important for the competition of the pathogen against host-associated bacteria (47,48). This suggests that variability in the expression of effector to compete against the plant microbiota might impact the pathogen’s ability to colonize the host, therefore, they could be important for the evolution of host specificity. The virulent and avirulent *Z. tritici* isolates showed differences in the intensity of the interactions, particularly in the growth enhancement of bacteria. We explored the underlying mechanism of growth enhancement of *Pseudomonas* bacteria and found that the growth enhancement of bacteria correlates with the hydrolysis of sucrose in the media by the virulent Zt469, but not in the avirulent Zt549 (Figure 6). This finding suggests variation in the expression of the pathogen invertase enzymes in the presence of competitive bacteria.

Positive and negative correlations between microorganisms can reflect direct and indirect interactions between microbial members (49). Under this assumption, the different *Z. tritici* isolates correlated with different microbial OTUs indicate their ability to establish specific direct interactions with them in the apoplast and/or on the leaf surface. However, we could not validate the majority of the correlations due to the difficulty of isolating the majority of the bacteria (67 of 2546 OTUs) and fungi (17 of 195 OTUs) associated with the plants. Particularly, our network predictions show that the virulent Zt469 correlates positively with more taxa than Zt549. This finding is in agreement with Zt469 inducing bigger enhancement halos in *A. cylindrica-*associated bacteria than Zt549. Nevertheless, *Z. passerinii* also shows growth enhancement of similar bacterial taxa (eg, Rodococcus and Streptomyces), suggesting that this activity might be common in *Zymoseptoria* species, but its relevance might also depend on the host context and what bacteria are prevalent in the microbiome.

### Bacterial growth enhancement correlates with sucrose hydrolysis in the media

Our data shows that the *Z. tritici* (Zt469, Zt549) and *Z. passerinii* (Zpa21) can hydrolyze sucrose in the media resulting in the diffusion of monosaccharides (Figure 6). These enzymes hydrolyze the glycosidic bond in sucrose, inulin, and levan and are present in plant-associated fungi to use plant-derived sugars, including fungal pathogens (50). Our experimental data suggests that the degradation of sucrose to monosaccharides by *Zymoseptoria* is the mechanism by which the pathogen enhances the growth of bacteria *in vitro*, pointing towards a substrate cross feeding model of interaction (51) between pathogen and leaf-associated bacteria.

Our genome analysis shows that Zt469, Zt549, Zpa796, and Zpa21 also possess invertase genes that are orthologous of the ones identified in *Z. tritici I*PO323 (Supplementary Table 10). However, only Zt469, Zt549, and Zpa21 show *in vitro* invertase activity in the media, while Zpa796 does not. Interestingly, only the isolate Zt469 shows invertase activity in the presence of *A. cylindrica-*associated *Pseudomonas*, which correlates with the observed growth enhancement of the bacteria. So far, the role of invertases in plant pathogens has been associated with their ability to consume the host’s sucrose (52,53), However, it has been suggested that *Zymoseptoria* relies on other diverse carbon sources, different from sucrose (36). Our work indicated that the activity of this enzyme might be relevant for the interaction of *Zymoseptoria* with the microbiome of its host. A detailed functional validation of these invertases will unravel their relevance in *Zymoseptoria* virulence and microbial interaction.

Pathogen-bacteria interactions with positive outcomes for the partners have so far not been studied much in the context of plant microbiomes (4). Our screening *in vitro* shows a possible mechanism by which bacteria can benefit from the presence of a fungal pathogen; however, *in vitro* interactions do not necessarily resemble what happens in planta, and the fitness outcomes might be different in the host context. Particularly, we find that many bacterial isolates that show growth enhancement around the pathogen also show antimicrobial activity against it (e.g. *Pseudomonas* Ac26, ac9, ac30, Supplementary Table 2). In bacteria, the synthesis of antimicrobials can be regulated by catabolite repression by sugars (54), and we speculate that *Zymoseptoria* could alleviate the competition of bacteria by modifying the sugar dynamics in the host. Future research will focus on evaluating the significance of this interaction mechanism for the pathogen and microbiome in the host

Our study suggests that the leaf microbiome of wild grasses is affected by the biotrophic infection of *Zymoseptoria* pathogens. Interestingly, plant-associated fungi suffer a higher loss of diversity than bacteria during pathogen invasion. Detailed community analyses suggest that virulent and avirulent *Zymoseptoria* isolates correlate with different microbial OTUs. Despite the difficulty of separating direct and indirect effects of fungal invasion on the phyllosphere microbiome composition, we conclude that *Zymoseptoria* can inhibit and enhance the growth of certain bacteria *in vitro*, suggesting that such direct effects also occur *in planta*. The virulent *Z. tritic*i and *Z. passerinii* isolates show similar phenotypes when confronted with the microbiome of their respective host. However, only Zt469 enhances the growth of *A. cylindrica-*associated *Pseudomonas* through the activity of the sucrose-degrading enzymes, suggesting that this mechanism could be important for host specificity. Our data supports a scenario where pathogens establish interactions with host-microbiomes which are not only antagonistic or neutral, but also beneficial. The mechanisms underlying these interactions may explain the ability of pathogens to coexist in the same host niche.

## Author contribution

EHS: Funding acquisition and resources

VFN, MH, TDS, GR, SB: Performance of the research

VFN, EHS: Design of research, interpretation of the results and write the manuscript

VFN, EHS, MH, TDS, GR, SB: Review the manuscript

## Data availability

Raw amplicon sequencing data are available in the NCBI bioproject PRJNA1263788. The partial 16S and ITS from microbial isolates are available in the accession numbers PV697047 - PV697222 and PV702891- PV702932, respectively. Data and code for analysis are available in https://doi.org/10.5281/zenodo.15698536

## Acknowledgments

This work was funded by the European Research Council (ERC consolidator grant 101087809 FungalSecrets). We thank the Leibniz-Institut für Pflanzengenetik und Kulturpflanzenforschung (IPK Genebank, Leibniz Institute) for facilitating the wild barley accession. We thank the Environmental Genomics group for the methods and result discussions.

## References

1. Trivedi P, Leach JE, Tringe SG, Sa T, Singh BK. Plant–microbiome interactions: from community assembly to plant health. Nat Rev Microbiol. 2020;18(11):607–21.

2. Dastogeer K, Tumpa F, Sultana A, Akter M, Chakraborty A. Plant microbiome–an account of the factors that shape community composition and diversity. Curr Plant Biol 23: 100161. 2020.

3. Fitzpatrick CR, Salas-González I, Conway JM, Finkel OM, Gilbert S, Russ D, et al. The plant microbiome: from ecology to reductionism and beyond. Annu Rev Microbiol. 2020;74(1):81–100.

4. Flores-Nunez VM, Stukenbrock EH. The impact of filamentous plant pathogens on the host microbiota. BMC Biol. 2024;22(1):175.

5. Fagundes WC, Hansen R, Rojas Barrera IC, Caliebe F, Feurtey A, Haueisen J, et al. Host specialization defines the emergence of new fungal plant pathogen populations [Internet]. Evolutionary Biology; 2024 [cited 2025 Jan 28]. Available from: http://biorxiv.org/lookup/doi/10.1101/2024.09.30.615799

6. Haueisen J, Stukenbrock EH. Life cycle specialization of filamentous pathogens — colonization and reproduction in plant tissues. Curr Opin Microbiol. 2016 Aug;32:31–7.

7. Snelders NC, Rovenich H, Petti GC, Rocafort M, van den Berg GCM, Vorholt JA, et al. Microbiome manipulation by a soil-borne fungal plant pathogen using effector proteins. Nat Plants. 2020 Nov;6(11):1365–74.

8. Snelders NC, Boshoven JC, Song Y, Schmitz N, Fiorin GL, Rovenich H, et al. A highly polymorphic effector protein promotes fungal virulence through suppression of plant-associated Actinobacteria. New Phytol. 2023;237(3):944–58.

9. Snelders NC, Petti GC, van den Berg GCM, Seidl MF, Thomma BPHJ. An ancient antimicrobial protein co-opted by a fungal plant pathogen for in planta mycobiome manipulation. Proc Natl Acad Sci. 2021 Dec 7;118(49):e2110968118.

10. Snelders NC, Rovenich H, Thomma BP. Microbiota manipulation through the secretion of effector proteins is fundamental to the wealth of lifestyles in the fungal kingdom. FEMS Microbiol Rev. 2022;46(5):fuac022.

11. Soldan R, Fusi M, Cardinale M, Daffonchio D, Preston GM. The effect of plant domestication on host control of the microbiota. Commun Biol. 2021;4(1):936.

12. Abdullaeva Y, Ratering S, Ambika Manirajan B, Rosado-Porto D, Schnell S, Cardinale M. Domestication Impacts the Wheat-Associated Microbiota and the Rhizosphere Colonization by Seed- and Soil-Originated Microbiomes, Across Different Fields. Front Plant Sci. 2022 Jan 12;12:806915.

13. Gholizadeh S, Mohammadi SA, Salekdeh GH. Changes in root microbiome during wheat evolution. BMC Microbiol. 2022 Dec;22(1):64.

14. Gruet C, Muller D, Moënne-Loccoz Y. Significance of the Diversification of Wheat Species for the Assembly and Functioning of the Root-Associated Microbiome. Front Microbiol. 2022 Jan 4;12:782135.

15. Bulgarelli D, Garrido-Oter R, Münch PC, Weiman A, Dröge J, Pan Y, et al. Structure and function of the bacterial root microbiota in wild and domesticated barley. Cell Host Microbe. 2015;17(3):392–403.

16. Escudero-Martinez C, Coulter M, Alegria Terrazas R, Foito A, Kapadia R, Pietrangelo L, et al. Identifying plant genes shaping microbiota composition in the barley rhizosphere. Nat Commun. 2022;13(1):3443.

17. Hassani MA, Özkurt E, Franzenburg S, Stukenbrock EH. Ecological assembly processes of the bacterial and fungal microbiota of wild and domesticated wheat species. Phytobiomes J. 2020;4(3):217–24.

18. Özkurt E, Hassani MA, Sesiz U, Künzel S, Dagan T, Özkan H, et al. Seed-Derived Microbial Colonization of Wild Emmer and Domesticated Bread Wheat (*Triticum dicoccoides* and *T. aestivum*) Seedlings Shows Pronounced Differences in Overall Diversity and Composition. Di Pietro A, editor. mBio. 2020 Dec 22;11(6):e02637–20.

19. Wipf HML, Coleman-Derr D. Evaluating domestication and ploidy effects on the assembly of the wheat bacterial microbiome. Zhang A, editor. PLOS ONE. 2021 Mar 18;16(3):e0248030.

20. Abdullaeva Y, Ratering S, Rosado-Porto D, Manirajan BA, Glatt A, Schnell S, et al. Domestication caused taxonomical and functional shifts in the wheat rhizosphere microbiota, and weakened the natural bacterial biocontrol against fungal pathogens. Microbiol Res. 2024;281:127601.

21. Deng L, Zhang A, Wang A, Zhang H, Wang T, Song W, et al. Wheat domestication alters root metabolic functions to drive the assembly of endophytic bacteria. Plant J. 2024 Nov;120(4):1263–77.

22. Rojas Barrera IC, Fagundes WC, Stukenbrock EH. Species of Zymoseptoria (Dothideomycetes) as a Model System to Study Plant Pathogen Genome Evolution. In: Plant Relationships: Fungal-Plant Interactions. Springer; 2022. p. 349–70.

23. Rojas-Barrera IC, Flores-Núñez VM, Haueisen J, Alizadeh A, Salimi F, Stukenbrock EH. Evolution of sympatric host-specialized lineages of the fungal plant pathogen Zymoseptoria passerinii in natural ecosystems. New Phytol. 2025;245(4):1673–87.

24. Haueisen J, Möller M, Seybold H, Small C, Wilkens M, Jahneke L, et al. Comparative analyses of compatible and incompatible host-pathogen interactions provide insight into divergent host specialization of closely related pathogens. Mol Plant Microbe Interact. 2025;MPMI–10.

25. Haueisen J, Möller M, Eschenbrenner CJ, Grandaubert J, Seybold H, Adamiak H, et al. Highly flexible infection programs in a specialized wheat pathogen. Ecol Evol. 2019 Jan;9(1):275–94.

26. Seybold H, Demetrowitsch TJ, Hassani MA, Szymczak S, Reim E, Haueisen J, et al. A fungal pathogen induces systemic susceptibility and systemic shifts in wheat metabolome and microbiome composition. Nat Commun. 2020 Dec;11(1):1910.

27. Ware SB, Verstappen ECP, Breeden J, Cavaletto JR, Goodwin SB, Waalwijk C, et al. Discovery of a functional Mycosphaerella teleomorph in the presumed asexual barley pathogen Septoria passerinii. Fungal Genet Biol. 2007 May;44(5):389–97.

28. Fagundes WC, Haueisen J, Stukenbrock EH. Dissecting the Biology of the Fungal Wheat Pathogen *Zymoseptoria tritici* : A Laboratory Workflow. Curr Protoc Microbiol [Internet]. 2020 Dec [cited 2022 Jul 19];59(1). Available from: https://onlinelibrary.wiley.com/doi/10.1002/cpmc.128

29. Flores-Núñez VM, Camarena-Pozos DA, Chávez-González JD, Alcalde-Vázquez R, Vázquez-Sánchez MN, Hernández-Melgar AG, et al. Synthetic communities increase microbial diversity and productivity of Agave tequilana plants in the field. Phytobiomes J. 2023;7(4):435–48.

30. R Core Team. R: A Language and Environment for Statistical Computing [Internet]. Vienna, Austria: R Foundation for Statistical Computing; 2021. Available from: https://www.R-project.org/

31. Wickham H. ggplot2: Elegant Graphics for Data Analysis [Internet]. Springer-Verlag New York; 2016. Available from: https://ggplot2.tidyverse.org

32. Lyda TA, Joshi MB, Andersen JF, Kelada AY, Owings JP, Bates PA, et al. A unique, highly conserved secretory invertase is differentially expressed by promastigote developmental forms of all species of the human pathogen, Leishmania. Mol Cell Biochem. 2015;404:53–77.

33. Emms DM, Kelly S. OrthoFinder: solving fundamental biases in whole genome comparisons dramatically improves orthogroup inference accuracy. Genome Biol. 2015;16:1–14.

34. Buchfink B, Xie C, Huson DH. Fast and sensitive protein alignment using DIAMOND. Nat Methods. 2015;12(1):59–60.

35. Teufel F, Almagro Armenteros JJ, Johansen AR, Gíslason MH, Pihl SI, Tsirigos KD, et al. SignalP 6.0 predicts all five types of signal peptides using protein language models. Nat Biotechnol. 2022;40(7):1023–5.

36. Rudd JJ, Kanyuka K, Hassani-Pak K, Derbyshire M, Andongabo A, Devonshire J, et al. Transcriptome and metabolite profiling of the infection cycle of Zymoseptoria tritici on wheat reveals a biphasic interaction with plant immunity involving differential pathogen chromosomal contributions and a variation on the hemibiotrophic lifestyle definition. Plant Physiol. 2015;167(3):1158–85.

37. Quiza L, Tremblay J, Pagé AP, Greer CW, Pozniak CJ, Li R, et al. The effect of wheat genotype on the microbiome is more evident in roots and varies through time. ISME Commun. 2023;3(1):32.

38. Dilla-Ermita CJ, Lewis RW, Sullivan TS, Hulbert SH. Wheat genotype-specific recruitment of rhizosphere bacterial microbiota under controlled environments. Front Plant Sci. 2021;12:718264.

39. Agler MT, Ruhe J, Kroll S, Morhenn C, Kim ST, Weigel D, et al. Microbial Hub Taxa Link Host and Abiotic Factors to Plant Microbiome Variation. PLoS Biol. 2016 Jan;14(1):e1002352.

40. Hansen R, Fagundes WC, Stukenbrock EH. Comparative Transcriptomic and Microscopic Analyses of a Wild Wheat Relative Reveal Novel Mechanisms of Immune Suppression by the Pathogen Zymoseptoria tritici. Mol Plant Microbe Interact. 2025;(ja).

41. Sapkota R, Jørgensen LN, Boeglin L, Nicolaisen M. Fungal communities of spring barley from seedling emergence to harvest during a severe Puccinia hordei epidemic. Microb Ecol. 2023;85(2):617–27.

42. Kettles GJ, Bayon C, Sparks CA, Canning G, Kanyuka K, Rudd JJ. Characterization of an antimicrobial and phytotoxic ribonuclease secreted by the fungal wheat pathogen Zymoseptoria tritici. New Phytol. 2018;217(1):320–31.

43. de Guillen K, Mammri L, Gracy J, Padilla A, Barthe P, Hoh F, et al. Zymoseptoria tritici proteins structurally related to UmV-KP4 and UmV-KP6 are toxic to fungi, and define novel structural families of fungal effectors. 2024;

44. Fagundes WC, Moeller M, Feurtey A, Hansen R, Haueisen J, Salimi F, et al. Interspecies hybridization as a route of accessory chromosome origin in fungal species infecting wild grasses. bioRxiv. 2024;2024–10.

45. Yang R, Du X, Khojasteh M, Shah SMA, Peng Y, Zhu Z, et al. Green guardians: The biocontrol potential of Pseudomonas-derived metabolites for sustainable agriculture. Biol Control. 2025;105699.

46. Sharma V, Salwan R, Sharma PN, Kanwar S. Elucidation of biocontrol mechanisms of Trichoderma harzianum against different plant fungal pathogens: Universal yet host specific response. Int J Biol Macromol. 2017;95:72–9.

47. Ökmen B, Katzy P, Huang L, Wemhöner R, Doehlemann G. A conserved extracellular Ribo1 with broad-spectrum cytotoxic activity enables smut fungi to compete with host-associated bacteria. New Phytol. 2023;240(5):1976–89.

48. Wang C, Han M, Min Y, Hu J, Pan Y, Huang L, et al. Colletotrichum fructicola co-opts cytotoxic ribonucleases that antagonize host competitive microorganisms to promote infection. Mbio. 2024;15(8):e01053–24.

49. Lee KK, Kim H, Lee YH. Cross-kingdom co-occurrence networks in the plant microbiome: Importance and ecological interpretations. Front Microbiol. 2022;13:953300.

50. Parrent JL, James TY, Vasaitis R, Taylor AF. Friend or foe? Evolutionary history of glycoside hydrolase family 32 genes encoding for sucrolytic activity in fungi and its implications for plant-fungal symbioses. BMC Evol Biol. 2009;9:1–16.

51. Wang C, Kuzyakov Y. Mechanisms and implications of bacterial–fungal competition for soil resources. ISME J. 2024;18(1):wrae073.

52. Chang Q, Liu J, Lin X, Hu S, Yang Y, Li D, et al. A unique invertase is important for sugar absorption of an obligate biotrophic pathogen during infection. New Phytol. 2017;215(4):1548–61.

53. Kagda MS, Martínez-Soto D, Ah-Fong AM, Judelson HS. Invertases in Phytophthora infestans localize to haustoria and are programmed for infection-specific expression. MBio. 2020;11(5):10–1128.

54. Liu YH, Song YH, Ruan YL. Sugar conundrum in plant–pathogen interactions: roles of invertase and sugar transporters depend on pathosystems. J Exp Bot. 2022;73(7):1910–25.

