## Supplementary Material for "Host-specific fungal plant pathogens exhibit distinct interactions with the leaf microbiota of wild grasses"

### Supplementary Methods

#### Culture conditions

*Zymoseptoria* inocula were obtained by cultivating the preserved isolates (glycerol 40% at -80°C) in YMS broth (yeast extract 4 g/L, malt extract 4g/L, sucrose 4g/L, pH=7.2) at 18°C, 200 rpm, for 5 days (Supplementary Table 1). Cells were harvested by centrifugation (10,000 rpm, 10 min), the supernatant was removed, and the cells were resuspended in either sterile PBS 1X (pH=7.4, Roth, Art.-Nr. 1108.1) or sterile millipore water.

#### Plant propagation and infection assays

*A. cylindrica* and *H. murinum* seeds were sown in a 729 ml pot filled with peat substrate inoculated with a soil slurry (1g of sifted soil in 50 ml of sterile distilled water) made with soil from the experimental farm of the Christian-Albrechts University Kiel "Hohenschulen" (Kiel, Germany). After germination, only one seedling per pot was maintained under greenhouse conditions (~60-70% humidity, ~20°C/day, ~12°C/night, and a day/night cycle of 16h/8h) with three times of watering per week.  *A. cylindrical* and *H. murinum* plants were inoculated with spores of *Zymoseptoria* after 21 and 14 days post-sowing, respectively.

Plant infection was performed following the protocol described in Fagundes et al. (1) with some modifications. In brief, suspensions of the virulent and avirulent *Zymoseptoria* cells were adjusted to 0.2 OD (optical density at 600 nm) in sterile Tween-20 0.1% (v/v in sterile millipore water). Each suspension was inoculated onto the adaxial side of the second unfolded leaf (the first after the cotyledon leaf) of each seedling with an ethanol-disinfected spray gun (2.0 bar). Mock control plants were treated with 0.1% Tween-20, also using a spray gun. After the inoculum had dried on the leaf surface, the seedlings were enclosed in a plastic bag with 2 L of tap water to create a high-humidity environment to facilitate infection. After 2 days, plants were taken out of the bag and randomized in trays under the greenhouse conditions described above.

**Confocal microscopy**

Additionally, a 1 cm piece of each sampled leaf was destained in 2 ml of ethanol 99% to evaluate the *Zymoseptoria* infection using confocal microscopy following the protocol described in Rojas-Barrera et al. (2). Five leaf pieces per treatment, host, and time were imaged.

**DNA extraction, library preparation, and amplicon sequencing**

The metagenomic DNA from washed leaf samples was extracted with the FastDNA Spin Kit for Soil (MP Biomedicals, Santa Ana, USA). The mechanical lysis and DNA extraction were performed following the procedure described in Seybold et al. (3).

Library preparation and amplicon sequencing of the 16S-rRNA-V4 and ITS2 markers, for prokaryotic and eukaryotic microbial communities, respectively, were performed at the University of Minnesota Genome Center (Minneapolis, MN, USA). In brief, the metagenomic DNA was processed by creating 1:8 and 1:64 dilutions of each sample and subjecting them to qPCR using the primers V4_515F_Nextera and V4_806R_Nextera in the presence of 0.25 µM of PNA blockers (PNA Bio, Newbury Park, CA) targeting the V4 locus (95°C-5 min; 35 cycles of 98°C-0.20 sec, 75°C-10 sec, 55°C-15 sec, 72°C-1 min, and 72°C-5 min of final extension), and the primers ITS4_Nextera and 5.8SR_Nextera targeting the ITS2 locus (95°C-5 min; 35 cycles of 98°C-0.20 sec, 67.5°C-15 sec, 72°C-45 sec, and 72°C-5 min of final extension). Amplicons were generated by subjecting the sample to 25-cycle PCRs with the same primers and PCR conditions. PCR products were diluted 1:100 in water and subjected to a second 10-cycle PCR to attach the Illumina sequencing DNA regions and the sample's barcodes. Samples were uniquely dual-indexed, as detailed in Gohl, et al. (4). The PCR products were normalized using SequalPrep kits (Invitrogen), pooled into sequencing libraries, and cleaned with AMPure XP mag beads (Beckman Coulter). Sequencing libraries were loaded onto an Illumina MiSeq using a 2 x 300 v3 flow cell (Illumina, San Diego, CA). Additional sequencing of the 16S-rRNA-V5-V7 marker was performed following the protocol reported by Seybold et al. (3).

**Data processing**

The raw reads of 132 libraries were processed using the pipeline described in Flores-Nunez et al. (5) with some modifications. In brief, low-quality reads were trimmed and the primers were removed using Trimmomatic (6), then the trimmed reads were merged, filtered by quality and size, dereplicated, and clustered in Operational Taxonomic Units (OTU) using the VSEARCH pipeline (7). After removing chimeric sequences, the concatenated merged reads were mapped against the OTU sequences to generate an OTU table. Finally, the OTU sequences were classified using the SINTAX algorithm with the RDP (v19) database for the 16S-rRNA-V4 (8) and the Unite database (v9) for the ITS2 (9). Unclassified OTUs and those matching the host's genomic and mitochondrial DNA were removed from the OTU table. Furthermore, samples with less than 350 (16S-rRNA-V4) and 100 reads (ITS2) were removed, and OTUs with less than 2 reads or only present in one sample were removed. After filtering, 4,570,240 reads distributed across 116 samples were used for further analysis.

**Bacterial and fungal isolation**

To obtain a culture collection of bacterial and fungal endophytes that co-exist with *Zymoseptoria* pathogens, we established a culture collection of both bacteria and fungi from *A. cylindrica* and *H. murinum* leaves. To this end, we collected and washed leaf samples from the infection experiment. The washed leaves were ground with pestles in 3 ml of sterile PBS 1X in sterile mortars. The resulting cell suspensions were ten-folded diluted and 200 ul of each of the original suspension and dilutions were inoculated in agar plates containing Trypticase soy agar (TSA, Difco™, 211768, DB, France), Potato Dextrose agar (PDA, Difco™, 213400, DB, France), Czapek-Dox agar (CZ, Difco™, 233910, DB, France), or Grass-agar (40 g of blended leaves from *A. cylindrica* or *H. murinum* in 1 L of PBS 1X and 15 g/L of agar (Difco™, 214030, DB, France), pH = 7). All the plates were incubated at 23°C for 14 days. Bacterial and fungal colonies were picked at 3, 7, and 14 days post-inoculation, and subcultured in TSA and PDA, respectively. Colonies were classified in morphotypes based on morphological characteristics, and only 1 morphotype per treatment and time was further characterized. All the isolates were long-term stored in glycerol 40% at -80°C.

**Data processing and analysis**

All downstream and statistical analyses were performed in R (10), and all plots were constructed using the ggplot2 (11) package. First, the OTU counts were subsampled to account for differences in library size using the vegan package (12). Microbial diversity was analyzed between host, infection treatments, and time; and the differences between groups of samples were tested using a Kruskal-Wallis rank sum test with a Dunn *post hoc* test. Alpha diversity was evaluated by calculating the OTU richness (number of OTUs per sample) and the Shannon index (abundance and evenness of OTUs). Beta diversity was estimated using the Bray-Curtis dissimilarity of the cumulative sum scaled OTU table (CSS, metagenomeseq package, (13,14)) and analyzed in a non-metric multidimensional scaling ordination (NMDS) plot and a PERMANOVA analysis (vegan package(12)).

The core microbiome was defined between hosts by selecting the abundant (<0.01% of relative abundance) and prevalent OTUs (<50 and 20% of the samples for 16S-V4-rRNA and ITS, respectively) between the groups of samples using the phyloseq (15) and microbiome package (16). The intersection between the core microbiome of each host was calculated using the ggVennDiagram (17) and upsetR (18) packages.

Correlation networks were constructed for both hosts infected with the virulent and avirulent *Zymoseptoria* lineage at 4 and 7 dpi (biotrophic phase) to assess the community structure and the correlations of *Zymoseptoria* with other OTUs. The network analysis was performed as described in Flores-Nunez et al. (5) with modifications. In brief, the prokaryotic and eukaryotic OTU tables were merged, and the correlations between OTUs were calculated using the sparCC (19) method in the spiecEasi package (20). Only correlations with a weight > |0.6| and a p-value < 0.01 were used for constructing an undirected network with the igraph (21) package, plotted using the GGally (22)package their network metrics were calculated. A subnetwork composed only of *Zymoseptoria*-correlating OTUs correlations was extracted, and the proportions of positive and negative correlations were calculated.

To determine if the differences in network structure were due to the treatment and not due to random associations, the analyses described above were performed in random subsampled networks. For each host and treatment combination, the OTU table was sampled 100 times, with the replacement of 50% of OTUs and 8 samples. Additionally, Random networks comprising all the data sets (both virulent and avirulent treatment) were constructed with the same parameters to compare the proportion of negative and positive correlations expected independently from the treatment. The metrics described above were calculated for all randomized networks, and the differences were evaluated using a Kruskal-Wallis rank sum test with a Dunn *post hoc* test.

A differential abundance analysis was performed with ANCOMBC2 (23)to determine the differential OTUs between hosts and between treatments in a single host. The analysis was performed with the raw OTU table with a prevalence threshold of 50% and 20% for the prokaryotic and eukaryotic microbial communities, respectively. Only the OTUs with a corrected p-value < 0.05 and a logFC > |1| were considered to be differentially abundant.

**Characterization of bacterial and fungal isolates**

Fungi and bacteria were characterized using the partial 16S and ITS amplicons, respectively. Bacterial and yeast-like isolates were grown on 5 mL of TSA for 1-3 days at 23°C and 200 rpm. Then, 50 µL of liquid culture was mixed with 100 µL of lysis buffer (0.1% Triton-100 in TE (10 mM TrisHCl pH=8, 10 mM EDTA)) and incubated at 99°C for 10 min at 500 rpm in a thermoblock. The supernatant was recovered after 10 min of centrifugation at 13,000 rpm. A 1:100 dilution of the supernatant was used as the template for the partial 16S-rRNA sequencing using the 16S-341F and 16S-1193R primer pair.

Filamentous fungi were grown in Potato Dextrose Broth (PDB, P6685, Sigma Aldrich$^{R}$) for 2-5 days at 23°C and 200 rpm. Then, 50 µL of the liquid culture or pieces of mycelial pellets were mixed with 100 µL of lysis buffer and processed as above. A 1:100 dilution of the supernatant was used as the template for the partial ITS amplification using the ITS1 and ITS4 primers.

All amplicons were purified and sequenced by Eurofins (Eurofins Umwelt Nord GmbH, Kiel, Germany) using the same primers as for amplification. The genus of each isolate was identified by BLAST against the NCBI 16S-rRNA and ITS reference database.

To match the isolates to the amplicon sequencing data. The V4 and ITS2 region was extracted from the aligned 16S-rRNA and ITS partial sequences. Then, unique phylotypes were determined by clustering the extracted regions using the --cluster_size function of VSEARCH at 100% sequence identity. Finally, the unique phylotypes were mapped against the OTU sequences using the --usearch_global VSEARCH function at 97% and 95% of sequence identity for the 16S-rRNA-V4 and ITS2 data, respectively.

**Confrontation assays**

The bacterial isolates and four *Zymoseptoria* isolates were confronted in two one-way assays on Fries-3 media 0.5X (2.5 g/L C4H12N2O6, 0.5 g/L NH4NO3, 0.25 g/L, MgSO4*7H2O, 0.65 g/L KH2PO4, 1.3 g/L K2HPO4, 0.5 g/L yeast extract, 15 g/L sucrose, 7.5 g/L agar) supplemented with 1 mL of trace elements (167 mg/L LiCl, 227 mg/L CuSO4.5H2O, 34 mg/L H2MoO4, 72 mg/L MnCl2.4H2O, 80 mg/L470, CoCl2.6H2O).

For the first assay, 15 µL of every bacterial suspension of 0.2 of optical density (OD, 600 nm) was mixed with 15 ml of Fries-3 media 0.5X, maintained between 40-50°C and poured in a petri dish. After solidification, the bacteria-agar was divided into 4 sections, and a droplet of 1.5 ul of a *Zymoseptoria* suspension of 2.7 OD was added on top by triplicate. Additionally, a droplet of 1.5ul of gentamicin (25 mg/ml) was added as a positive control of inhibition in the fourth section. In the second assay, 1.5 ml of every *Zymoseptoria* suspension of 0.2 OD was mixed with 13.5 ml of Fries-3 media 0.5X maintained between 40-50°C and poured into a petri dish. After solidification, the plate was divided into 4 sections, and a droplet of 1.5 ul of a bacterial suspension of 0.2 OD was added on top of the bacteria-agar by triplicate. Additionally, 1.5ul of hygromycin (25 mg/ml) was added as a positive control of inhibition in the fourth section.

Every combination of bacteria and *Zymoseptoria* isolates was tested in an individual Petri dish. Bacteria-agar plates and *Zymoseptoria*-agar plates were incubated at 18°C and 23°C for 7 days, respectively, to give advantage to the microorganism growing on top. After incubation, the microorganism embedded in the agar grows as a monolayer, while the microorganism inoculated on the top grows as a single colony. When a top isolate grew surrounded by a clearance halo in the monolayer isolate, it was considered an inhibition phenotype (I). It was considered a growth enhancement phenotype (P) when a top isolate grew surrounded by a denser halo in the monolayer isolate. The absence of inhibition or enhancement halos in the monolayer isolate or the absence of growth in the top isolate was considered a non-effect phenotype (N). The diameter of the colony at the top and the diameter of the growth inhibition or enhancement halos were measured manually with a ruler.

We also tested the interaction between fungal species in an *in vitro* assay. The filamentous fungi and isolates of *Zymoseptoria* were used in a two-way confrontation assay. A 0.5 cm mycelium plug was inoculated in the center of an agar plate with 15 ml of Fries-3 0.5X, and 4 droplets of 1.5 µL of a *Zymoseptoria* suspension (2.7 OD) were inoculated in 4 equidistant points 2 cm apart from the mycelium plug. Plates were incubated at 23°C, and the radius of the filamentous fungal colony towards the pathogen was recorded together with the *Zymoseptoria* colony diameter. The outcome of the confrontation was evaluated by measuring the diameter of the colonies, growing together and alone, and calculating the log2 fold-change between them (lfc>0.42). For the endophytic yeasts, we assessed their interaction with *Zymoseptoria* as in the bacterial confrontation assay described above.

**Influence of pathogen concentration and sugar on *in vitro* confrontation assays**

The confrontation assays were performed as in the previous section with modifications. The *Zymoseptoria* OD was adjusted to 2.7 and 1.35, and the assays were performed in Fries-3 media 0.5x with sucrose or with glucose. Plates were incubated at 18°C for 7 days, and the presence and diameter of colonies and growth-enhancement halo were recorded.

Since the growth enhancement of bacteria did not happen in the presence of glucose, we evaluated the sucrose degradation activity in the confrontation assays with a 2,3,5-triphenyl tetrazolium chloride (TTC) staining. The protocol described in Lyda et al. (24) was followed with some modifications. In brief, 5 ml of 0.2% TTC in NaOH (1M) was gently poured into the agar and incubated for 10 min at room temperature. Finally, the TTC solution is removed from the petri dish. The formation of a red precipitate surrounding the *Zymoseptoria* colony indicates the presence of reducing hexoses (glucose and fructose) derived from the sucrose in the media. Additionally, a plate with bacteria embedded in the agar media with sterile paper discs with 10 uL of glucose 2 M was used as a positive control of growth enhancement.

**Orthology inference**

Orthologous relationships among proteins from Zymoseptoria species were inferred using OrthoFinder v2.2.7(25), implementing DIAMOND v0.9.24.125 (26) for the all-versus-all comparisons step. The analysis included proteomes from the following species: *Z. tritici*, isolates IPO323 (27), Zt469 (1), and Zt549 (BioSample SAMN43819765, 1); *Z. passerinii*, isolates Zpa796 (2) and  Zpa63 (28); Z. ardabiliae Za17, Z. brevis Zb87, and Z. pseudotritici Zp13 (28); Z. crescenta CBS 144410 (BioSample SAMN27594421); and Z. verkleyi Zv3 (Tanneau et al., unpublished data). The proteome of Cercospora beticola Cb09-40 (29) was used as outgroup.

**Genome assembly and gene prediction**

Gene models and assembled genome data was not available for Z. crescenta CBS 144410 and Z. tritici Zt549, respectively. Thus, we performed de novo genome assembly for Zt549 using the Illumina reads from NCBI. Read quality was assessed using FastQC v0.11.9 (https://www.bioinformatics.babraham.ac.uk/projects/fastqc/). Adapter trimming and quality filtering were performed with Cutadapt v3.7 (30), using the parameters --minimum-length=50, --max-n=2, and --quality-cutoff=30. Genome assembly was carried out with SPAdes v3.15.0 (31) using trimmed reads and the parameters “-k 21,33,55,67,99,127” and “--careful”. Contigs shorter than 200 bp were removed using seqtk v1.4-r122 with the flag “-L 200”. Assembly quality was evaluated with QUAST v5.0.2 (32), using the parameters --fungus and --fragmented. Completeness was assessed using BUSCO v5.8.2 (33) against the dothideomycetes_odb10 dataset, which includes 3,786 conserved fungal orthologs. Gene models for Z. tritici Zt549 and Z. crescenta CBS 144410 were predicted using BRAKER2 v2.1.6 (34), with parameters set to --alternatives-from-evidence=false. Protein homology evidence was supplied to using a custom protein database composed of fungal proteins retrieved from OrthoDB v12 (http://bioinf.uni-greifswald.de/bioinf/partitioned_odb12/). Additionally, species-specific protein hints were provided: Z. passerinii Zpa63 for Z. crescenta CBS 144410 and Z. passerinii IPO323 for Zt549.

### Supplementary Results

#### Core analysis

We characterized the bacterial core community of *A. cylindrica* and *H. murinum* mock-inoculated (uninfected) leaves by selecting the OTUs with a relative abundance >0.1% and a frequency of >50% in the samples. There was a core group of 26 and 19 bacterial OTUs in *A. cylindrica* and *H. murinum* that accounted for 48.7% and 60.6% of the total relative abundance, respectively. The two hosts shared 12 core OTUs (Figure 1C), including *Ralstonia* (OTU_40) and *Paraburkholderia* (OTU_41) as the most abundant core OTUs (Supplementary Table 3). The rest of the core OTUs were unique to one of the two plant species (14 for *A. cylindrica* and 7 for *H. murinum,* Figure 1C), some of the core unique OTUs of *A. cylindrica* were not detectable in *H. murinum* such as *Vogesella* (OTU_212) or they were differentially abundant between the hosts like the group *Planctomycetia* (family *Isosphaeraceae*, Supplementary Table 3 and 4).

We also characterized the core fungal communities of *A. cylindrica* and *H. murinum* as we did for bacteria. Only 12 core OTUs were detected in *A. cylindrica* and 6 in *H. murinum* which represent a mean of 52.2% and 32.8% of the total relative abundance. Five core OTUs were shared between the two grass species  (Figure 1G) including *Pseudogymnoascus* (OTU_6), *Penicillium* (OTU _209), and 3 unidentified genera (Supplementary Table 3). Only 7 and 1 core OTUs were unique to *A. cylindrica* and *H. murinum,* respectively (Figure 1G). Differential OTUs between host included *Vishniacozyma* (OTU_17, not a member of the core fungal community) in *A. cylindrica* and *Penicillium* OTU_209 (core member) and OTU_25 (not core member, Supplementary table 5)

#### Microscopy

In both host, we observed *Zymoseptoria* blastospores and hyphae on the leaf cuticle. However, only the virulent isolates were able to colonize the mesophyll tissue asymptomatically to produce symptoms of disease and pycnidia, while the avirulent isolates were not able to invade the mesophyll. *Zymoseptoria* was not present in the mock-inoculated samples, confirming the absence of cross-infection in our experiment

The microscopy analyses were in support of the ITS2 barcode sequencing. The absolute abundance of *Zymoseptoria* reads in the samples reflected the fungal biomass as it was higher in the virulent interaction compared to the avirulent and mock samples (Supplementary Figure 4).

#### Network metrics

Interestingly the average degree of connectivity, as a measure of network complexity, consistently decreased for A. *cylindrica* when infected with either the virulent or avirulent isolate of *Z. tritici* in comparison to the Mock treatment. However, the same patter was consistent in *H. murinum* only when evaluated with the V5-V7-rRNA amplicon and to a lesser extent with the V4-rRNA amplicon (Supplementary Figure 8).

By focusing on the common OTUs correlating with *Zymoseptoria*, we determine that the core OTUs *Paraburkholderia* (OTU_41), *Rhodococcus* (OTU_140), and *Nitrososphaera* (OTU_225) had positive correlations with the virulent Zt469 but negative correlations with the avirulent Zt549. No common correlations were found between the two *Z. passerinii* isolates, but the avirulent Zpa21 also correlates negatively with *Paraburkholderia* (OTU_41) Supplementary Table 8).

When the number of positive or negative correlations was normalized against the average number of correlations per OTU, the virulent isolate Zt469 had almost twice the number of positive correlations than the average OTU in the network (mean ratio =1.9, Supplementary Figure 8). A similar pattern was observed for the dataset based on the 16S-rRNA-V5-V7 region (Supplementary Figure 9).

**Culture collection**

The leaf-associated bacteria and fungi were isolated from uninfected and infected leaves between 0dpi and 7dpi. For *A. cylindrica*, we obtained 117 bacterial and 26 fungal isolates corresponding to 28 and 11 different genera respectively. For *H. murinum*, we isolated 69 bacterial isolates and 16 fungal isolates corresponding to 25 and 8 different genera, respectively.

We used PCR amplification to determine the genera of the different microbiome members using primers targeting the 16S-rRNA and ITS loci. To associate our isolates with the amplicon sequencing data, the V4 and ITS2 region was extracted from the partial 16S rRNA and ITS marker sequences, respectively, de-replicated into phylotypes with 100% sequence identity and clustered to the OTU sequences at 97% and 95% of identity, respectively. The bacteria isolated from both hosts accounted for an average of 15.1% and 6.7% of the OTU relative abundance across the *A. cylindrica* and *H. murinum* samples, respectively, while the fungal isolates accounted for an average of 10.1% and 4.5%, respectively. The identity and metadata of each isolate can be consulted in our data repository.

### Supplementary Tables

**Supplementary Table 1.** *Zymoseptoria* strains used in this work.

| Species name | Strain name | Host | Country of origin | Description |
|---|---|---|---|---|
| *Zymoseptoria tritici* | Zt469 | *Aegilops cylindrica* | Iran | Fagundes, et al., 2025 (1) |
| *Zymoseptoria tritici* | Zt549 | *Triticum aestivum* | Iran | Fagundes, et al., 2025 (1) |
| *Zymoseptoria passerinii* | Zpa796 | *Hordeum murinum* subs. glaucum | Iran | Rojas-Barrera, et al. 2025 (2) |
| *Zymoseptoria passerinii* | Zpa21* | *Hordeum vulgare* subs. vulgare | USA | Ware, et al., 2007 (28, 35) |

* P63 in reference

**Supplementary Table 2.** List of *Pseudomonas* spp. isolates from *Aegilops cylindrica* as identified from 16S amplicon data and from the culture collection.

| Isolate id | Genus | 16S phylotype | Bacteria's effect on Zt469 | Zt469's effect on bacteria | OTU match | Mean read count *A. cylindrica** | Mean read count *H. murinum** |
|---|---|---|---|---|---|---|---|
| ac9 | *Pseudomonas* | Pse_4 | Inhibition | Enhancement | OTU_81 | 21 | 0 |
| ac26 | *Pseudomonas* | Pse_5 | Inhibition | Enhancement | OTU_74 | 33 | 1 |
| ac30 | *Pseudomonas* | Pse_6 | Inhibition | Enhancement | OTU_698 | 30 | 14 |
| ac66 | *Pseudomonas* | Pse_2 | No effect | Enhancement | OTU_37 | 0 | 3 |

*Mean value from 21 and 22 samples for *A. cylindrica* and *H. murinum*, respectively

**Supplementary Table 3.** PERMANOVA analysis of the microbiome of uninfected (mock-inoculated) *A. cylindrica* and *H. murinum* leaves.

| factor | Bacteria (16S-rRNA-V4) | | | | | Fungi (ITS2) | | | | |
|---|---|---|---|---|---|---|---|---|---|---|
| | Df | SumOfSqs | R2 | F | Pr(>F) | Df | SumOfSqs | R2 | F | Pr(>F) |
| **host.id** | 1 | 0.806 | 0.065 | 2.965 | **0.001** | 1 | 0.718 | 0.051 | 2.145 | **0.002** |
| **time** | 4 | 1.474 | 0.119 | 1.355 | **0.001** | 4 | 1.937 | 0.137 | 1.447 | **0.003** |
| **host.id:time** | 4 | 1.155 | 0.093 | 1.063 | 0.229 | 4 | 1.445 | 0.102 | 1.080 | 0.263 |

**Supplementary Table 4.** Core bacterial and fungal OTUs from *A. cylindrica* and *H. murinum.*

| otu.id | domain | genus | intersection |
|---|---|---|---|
| OTU_40 | d:Bacteria | g:Ralstonia | Ac_Hm |
| OTU_41 | d:Bacteria | g:Paraburkholderia | Ac_Hm |
| OTU_56 | d:Bacteria | f:Comamonadaceae | Ac_Hm |
| OTU_98 | d:Bacteria | g:Paraburkholderia | Ac_Hm |
| OTU_140 | d:Bacteria | g:Rhodococcus | Ac_Hm |
| OTU_149 | d:Bacteria | g:Staphylococcus | Ac_Hm |
| OTU_170 | d:Bacteria | g:Bradyrhizobium | Ac_Hm |
| OTU_178 | d:Bacteria | g:Cutibacterium | Ac_Hm |
| OTU_206 | d:Bacteria | f:Bacillaceae | Ac_Hm |
| OTU_211 | d:Bacteria | g:Gp16 | Ac_Hm |
| OTU_225 | d:Archaea | g:Nitrososphaera | Ac_Hm |
| OTU_564 | d:Bacteria | g:Paraburkholderia | Ac_Hm |
| OTU_36 | d:Bacteria | g:Pseudomonas | Ac |
| OTU_63 | d:Bacteria | g:Limnobacter | Ac |
| OTU_67 | d:Bacteria | f:Microbacteriaceae | Ac |
| OTU_97 | d:Bacteria | g:Paeniglutamicibacter | Ac |
| OTU_115 | d:Bacteria | g:Acinetobacter | Ac |
| OTU_147 | d:Bacteria | g:Rheinheimera | Ac |
| OTU_155 | d:Bacteria | g:Streptomyces | Ac |
| OTU_166 | d:Bacteria | g:Nocardioides | Ac |
| OTU_212 | d:Bacteria | g:Vogesella | Ac |
| OTU_230 | d:Bacteria | f:Isosphaeraceae | Ac |
| OTU_248 | d:Bacteria | g:Gp2 | Ac |
| OTU_271 | d:Bacteria | p:Actinomycetota | Ac |
| OTU_304 | d:Bacteria | p:Actinomycetota | Ac |
| OTU_1753 | d:Bacteria | g:Mycobacterium | Ac |
| OTU_228 | d:Bacteria | p:Actinomycetota | Hm |
| OTU_247 | d:Bacteria | g:Solirubrobacter | Hm |
| OTU_264 | d:Bacteria | p:Bacillota | Hm |
| OTU_272 | d:Bacteria | p:Actinomycetota | Hm |
| OTU_338 | d:Bacteria | d:Bacteria | Hm |
| OTU_372 | d:Bacteria | d:Bacteria | Hm |
| OTU_7747 | d:Bacteria | g:Roseateles | Hm |
| OTU_4 | d:Fungi | d:Fungi | Ac_Hm |
| OTU_6 | d:Fungi | g:Pseudogymnoascus | Ac_Hm |
| OTU_20 | d:Fungi | d:Fungi | Ac_Hm |
| OTU_33 | d:Alveolata | p:Ciliophora | Ac_Hm |
| OTU_209 | d:Fungi | g:Penicillium | Ac_Hm |
| OTU_5 | d:Viridiplantae | g:Chlamydomonas | Ac |
| OTU_8 | d:Fungi | g:Apiotrichum | Ac |
| OTU_16 | d:Fungi | o:Hypocreales | Ac |
| OTU_29 | d:Fungi | g:Botrytis | Ac |
| OTU_46 | d:Fungi | g:Schizothecium | Ac |
| OTU_98 | d:Fungi | d:Fungi | Ac |
| OTU_199 | d:Fungi | g:Dissoconium | Ac |
| OTU_41 | d:Fungi | g:Botryotrichum | Hm |

**Supplementary Table 5.** Differential bacterial taxa from *A. cylindrica* and *H. murinum* determined by ANCOMBC2

| Taxon | logFC | p_unadjusted | p_adjusted | Abundant in |
|---|---|---|---|---|
| family:Isosphaeraceae | -1.29773 | 0.00059 | 0.01403 | *A. cylindrica* |
| genus:Paludisphaera | -0.99504 | 0.00061 | 0.01403 | *A. cylindrica* |
| class:Planctomycetia | 1.02836 | 0.00025 | 0.01403 | *H. murinum* |

**Supplementary Table 6.** Differential fungal OTUs from *A. cylindrica* and *H. murinum* determined by ANCOMBC2

| otu.id | genus | logFC | p_unadjusted | p_adjusted | Abundant in |
|---|---|---|---|---|---|
| OTU_17 | g:Vishniacozyma | -1.354 | 0.011 | 0.039 | *A. cylindrica* |
| OTU_209 | g:Penicillium | 1.835 | 0.001 | 0.009 | *H. murinum* |
| OTU_25 | g:Penicillium | 1.868 | 0.002 | 0.014 | *H. murinum* |

**Supplementary Table 7.** PERMANOVA analysis of the microbiome of *A. cylindrica* and *H. murinum* leaves inoculated with virulent and avirulent *Zymoseptoria* lineages

| Host | factor | Bacteria (16S-rRNA-V4) | | | | | Fungi (ITS2) | | | | |
|---|---|---|---|---|---|---|---|---|---|---|---|
| | | Df | SumOfSqs | R2 | F | Pr(>F) | Df | SumOfSqs | R2 | F | Pr(>F) |
| Ac | Time | 4 | 1.571 | 0.109 | 1.599 | **0.001** | 4 | 1.716 | 0.103 | 1.734 | **0.001** |
| | Treatment | 2 | 0.494 | 0.034 | 1.007 | 0.395 | 2 | 0.934 | 0.056 | 1.888 | **0.001** |
| | Time : Treatment | 6 | 1.568 | 0.109 | 1.065 | 0.186 | 6 | 1.665 | 0.100 | 1.122 | 0.110 |
| Hm | Time | 4 | 1.446 | 0.087 | 1.284 | **0.001** | 4 | 0.986 | 0.096 | 1.534 | **0.003** |
| | Treatment | 2 | 0.524 | 0.032 | 0.930 | 0.796 | 2 | 1.994 | 0.193 | 6.206 | **0.001** |
| | Time : Treatment | 6 | 1.654 | 0.100 | 0.979 | 0.631 | 6 | 1.068 | 0.104 | 1.108 | 0.209 |

**Supplementary Table 8.** Zymoseptoria - OTU correlations in networks

| domain | correlating OTU | genus | weight of correlation | treatment | host |
|---|---|---|---|---|---|
| Bacteria | OTU_781 | g:Marmoricola | -0.655808 | Zt549 | Aegilops.cylindrica |
| Bacteria | OTU_41 | g:Paraburkholderia | -0.707766 | Zt549 | Aegilops.cylindrica |
| Bacteria | OTU_309 | g:Rhodopseudomonas | -0.601958 | Zt549 | Aegilops.cylindrica |
| Archaea | OTU_225 | g:Nitrososphaera | -0.628063 | Zt549 | Aegilops.cylindrica |
| Bacteria | OTU_140 | g:Rhodococcus | -0.689967 | Zt549 | Aegilops.cylindrica |
| Bacteria | OTU_115 | g:Acinetobacter | -0.747736 | Zt549 | Aegilops.cylindrica |
| Fungi | FOTU_93 | c:Sordariomycetes | -0.661835 | Zt549 | Aegilops.cylindrica |
| Protist | FOTU_5 | g:Chlamydomonas | 0.697838 | Zt549 | Aegilops.cylindrica |
| Bacteria | OTU_7747 | g:Roseateles | 0.688066 | Zt469 | Aegilops.cylindrica |
| Bacteria | OTU_605 | p:Actinomycetota | 0.618709 | Zt469 | Aegilops.cylindrica |
| Bacteria | OTU_564 | g:Paraburkholderia | 0.648817 | Zt469 | Aegilops.cylindrica |
| Bacteria | OTU_41 | g:Paraburkholderia | 0.756142 | Zt469 | Aegilops.cylindrica |
| Bacteria | OTU_40 | g:Ralstonia | 0.625666 | Zt469 | Aegilops.cylindrica |
| Bacteria | OTU_309 | g:Rhodopseudomonas | -0.649472 | Zt469 | Aegilops.cylindrica |
| Bacteria | OTU_303 | g:Gp16 | 0.800643 | Zt469 | Aegilops.cylindrica |
| Bacteria | OTU_271 | p:Actinomycetota | 0.798530 | Zt469 | Aegilops.cylindrica |
| Bacteria | OTU_2618 | f:Rhizobiaceae | -0.608690 | Zt469 | Aegilops.cylindrica |
| Bacteria | OTU_257 | g:Rheinheimera | -0.660281 | Zt469 | Aegilops.cylindrica |
| Bacteria | OTU_230 | f:Isosphaeraceae | -0.654298 | Zt469 | Aegilops.cylindrica |
| Archaea | OTU_225 | g:Nitrososphaera | 0.736816 | Zt469 | Aegilops.cylindrica |
| Bacteria | OTU_224 | g:Dyadobacter | -0.727041 | Zt469 | Aegilops.cylindrica |
| Bacteria | OTU_170 | g:Bradyrhizobium | -0.611674 | Zt469 | Aegilops.cylindrica |
| Bacteria | OTU_140 | g:Rhodococcus | 0.684659 | Zt469 | Aegilops.cylindrica |
| Fungi | FOTU_75 | g:Phialemonium | -0.660153 | Zt469 | Aegilops.cylindrica |
| Bacteria | OTU_267 | g:Paludisphaera | -0.670620 | Zpa796 | Hordeum.murinum |
| Bacteria | OTU_264 | f:Bacillaceae | 0.689441 | Zpa796 | Hordeum.murinum |
| Bacteria | OTU_198 | g:Neobacillus | 0.675070 | Zpa796 | Hordeum.murinum |
| Bacteria | OTU_41 | g:Paraburkholderia | -0.851360 | Zpa21 | Hordeum.murinum |
| Bacteria | OTU_338 | d:Bacteria | 0.643153 | Zpa21 | Hordeum.murinum |
| Bacteria | OTU_329 | g:Gp6 | -0.671075 | Zpa21 | Hordeum.murinum |
| Fungi | FOTU_67 | g:Tausonia | 0.617902 | Zpa21 | Hordeum.murinum |
| Fungi | FOTU_4 | d:Fungi | 0.702852 | Zpa21 | Hordeum.murinum |
| Fungi | FOTU_209 | g:Penicillium | 0.767076 | Zpa21 | Hordeum.murinum |

**Supplementary Table 9.** P-value tables from Fisher's exact test on contingency tables for the inhibition, promotion or no effect phenotypes on the confrontation assay between fungi and bacteria with the virulent and avirulent pathogens.

| Pathogen species | Fungi's effect on the pathogen | Pathogen´s effect on fungi | Bacteria's effect on the pathogen | Pathogen´s effect on the bacteria. |
|---|---|---|---|---|
| ***Z. tritici*** | 0.033 | 1 | 0.893 | 0.809 |
| ***Z. passerinii*** | 0.394 | 1 | 1 | 0.929 |

**Supplementary Table 10.** Orthologous genes in Zymoseptoria species with protein domains annotated as beta-fructofuranosidase activity

| Orthogroup | ***Z. tritici* IPO323** | ***Z. tritici* Zt469** | ***Z. tritici* Zt549** | ***Z. passerinii* Zpa796** | ***Z. passerinii* Zpa21** |
|---|---|---|---|---|---|
| OG0006625 | jgi\|Zymtr1\|53519\|ZtIPO323_114920.1(sp) | Zt469_000011F_arrow_0418.t1 | Zt549_g7767.t1 | Zpa796_jg9216.t1 | Zpa63_unitig_003_0399.t1 (sp) |
| OG0007405 | jgi\|Zymtr1\|57100\|ZtIPO323_040240.1 | Zt469_000001F_arrow_0975.t1 | Zt549_g2717.t1 | Zpa796_jg1395.t1 | Zpa63_unitig_004_0388.t1 |
| OG0007450 | jgi\|Zymtr1\|61540\|ZtIPO323_084640.1 (sp) | Zt469_000007F_arrow_0507.t1 (sp) | Zt549_g4596.t1 (sp) | Zpa796_jg10277.t1 (sp) | Zpa63_unitig_027_0094.t1 (sp) |
| OG0009431 | jgi\|Zymtr1\|58180\|ZtIPO323_051040.1 (sp) | Zt469_000002F_arrow_0053.t1 (sp) | Zt549_g10112.t1 | Zpa796_jg2600.t1 (sp) | |

sp: predicted secretion signal.

### Supplementary Figures

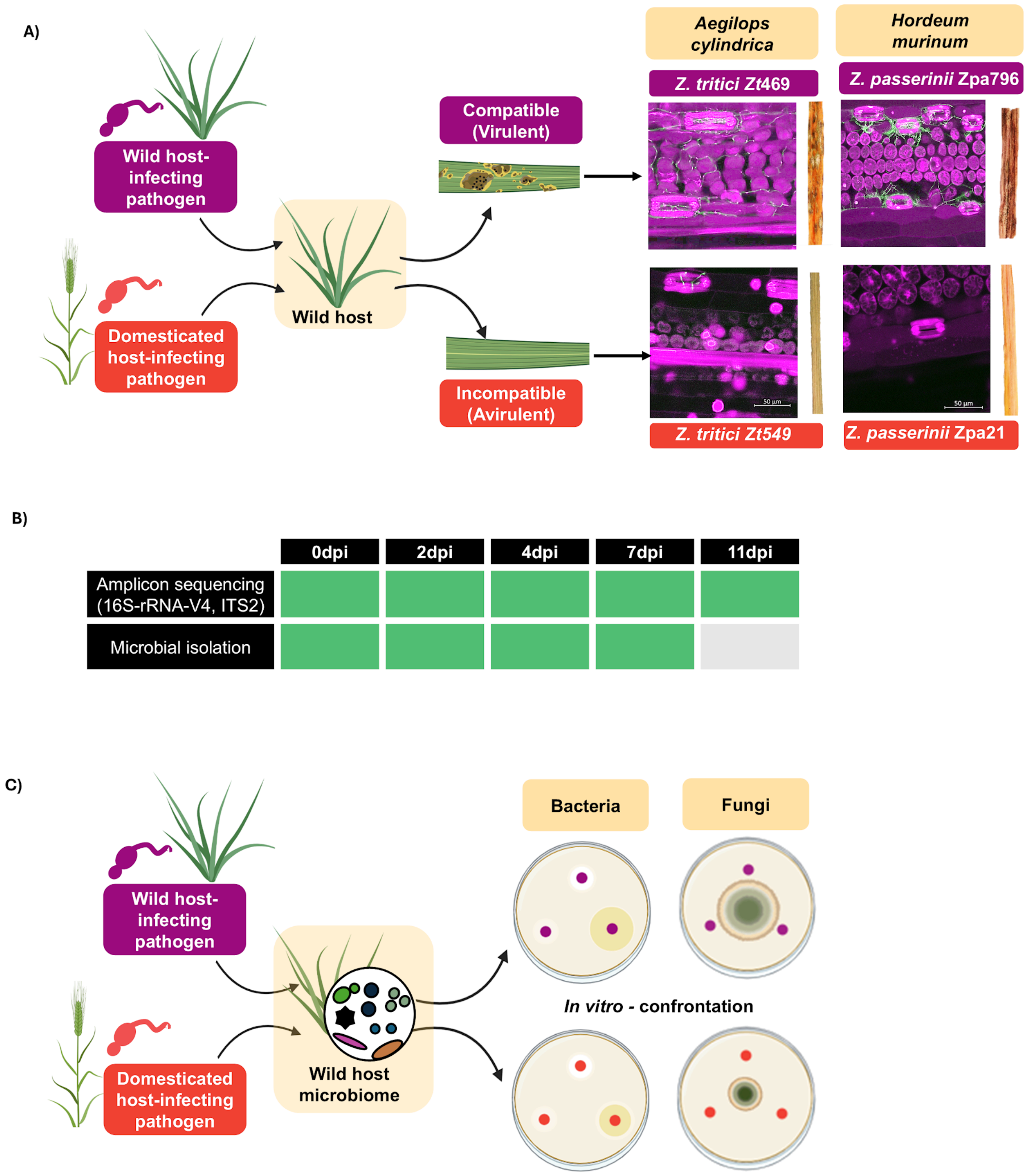


**Supplementary Figure 1.** Experimental design. A) The pathosystems used in this work. B) Sampling times for amplicon sequencing and microbial isolation. C) *In- vitro* confrontation test rationale.

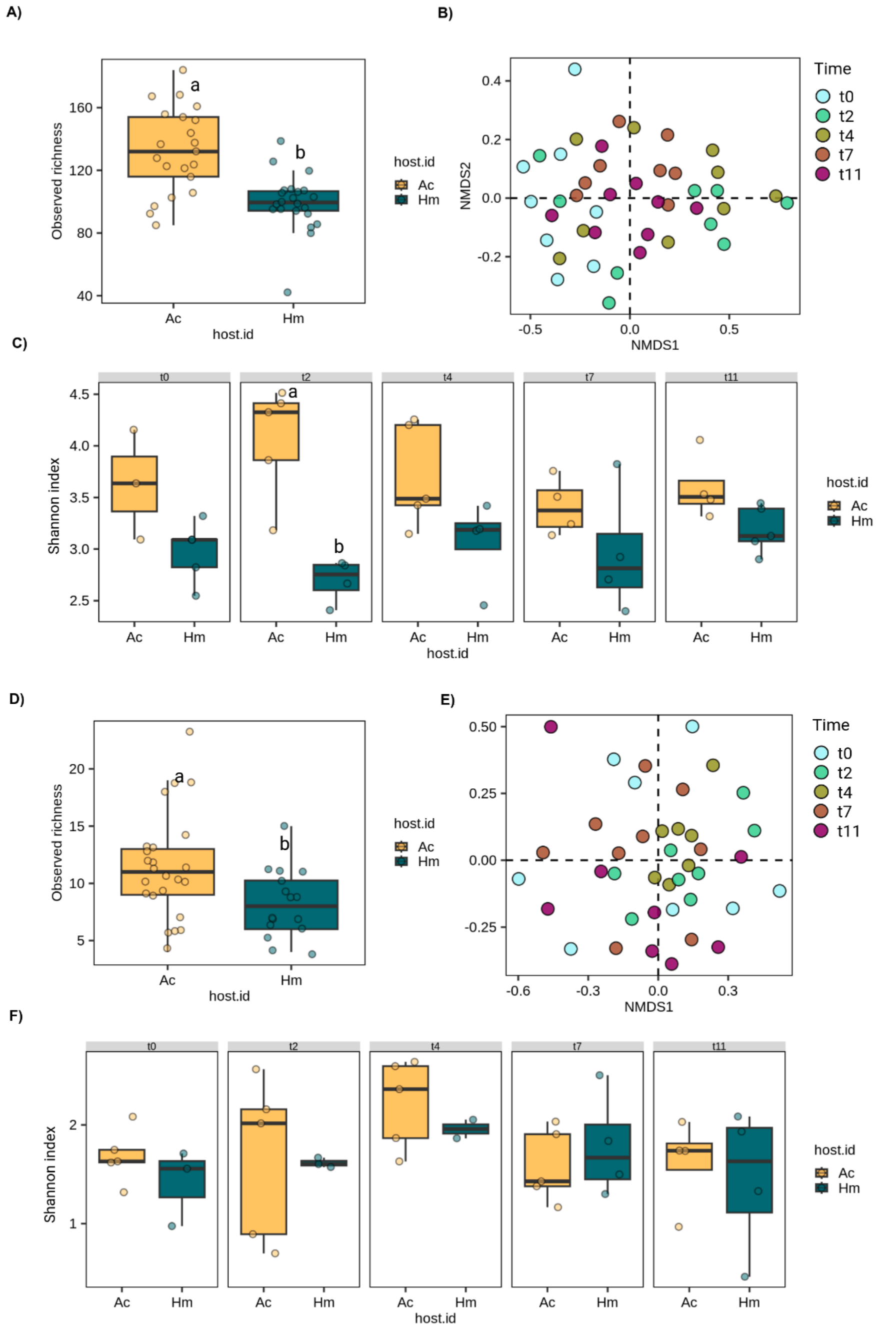


**Supplementary Figure 2**. The prokaryotic (A-C) and eukaryotic (D-F) leaf microbiome diversity of uninfected (mock-inoculated) leaves of *Aegilops cylindrica* (Ac) and *Hordeum murinum* (Hm) from 0 to 11 dpi. The panels show the OTU richness between host (A, D), the NMDS based on Bray-Curtis distances of the CSS normalized data between sampling times (B, E), and the Shannon index in each sampling point (C, F).

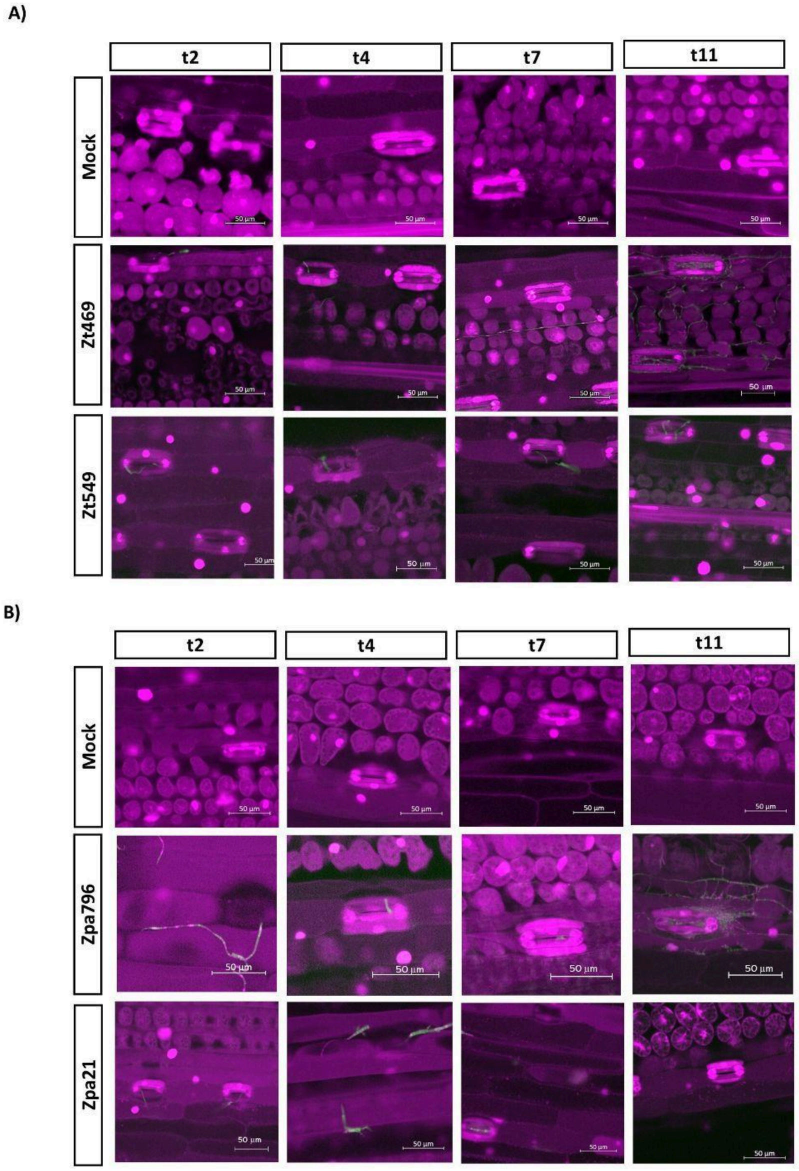


**Supplementary Figure 3**. Infection development of virulent *Z. tritici* Zt469 and avirulent *Z. tritici* Zt549 in *A. cylindrica* (A), and virulent *Z. passerinii* Zpa796 and avirulent *Z. passerinii* Zpa21 in *H. murinum* subs. *glaucum* (B).

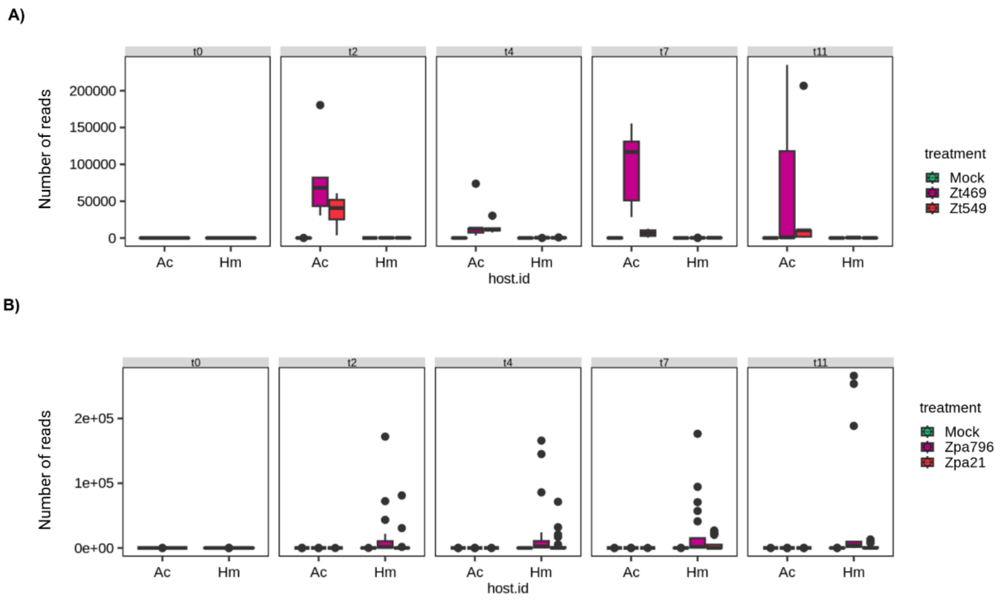


**Supplementary Figure 4**. Absolute read number of the OTUs classified as *Zymoseptoria* in the ITS2 amplicon sequencing data of *A. cylindrica* (A) and *H. murinum* subs. *glaucum* (B) infected with the virulent and avirulent strains, and the mock treatment.

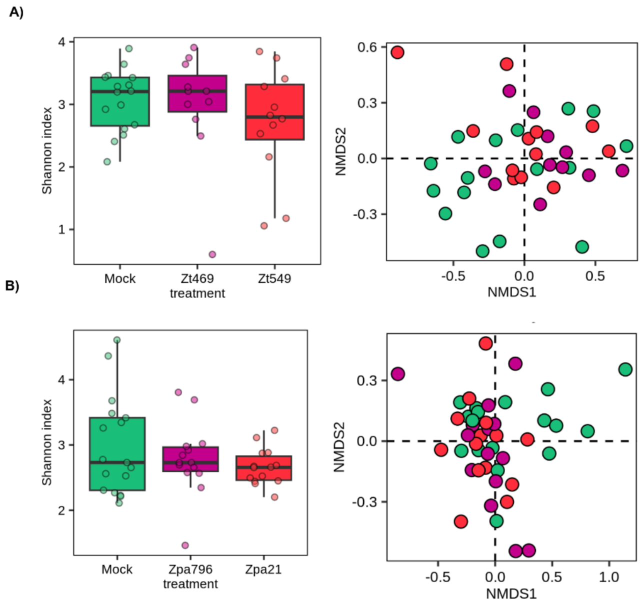


**Supplementary Figure 5**. Secondary 16S-rRNA-V5V7 metabarcoding of the prokaryotic microbiome of *A. cylindrica* (A) and *H. murinum* (B) infected with different lineages of *Z. tritici* and *Z. passerinii*, respectively. The boxplots represent the alpha diversity (Shannon index) in the mock, virulent, and avirulent treatments from all the time points while the NMDS were calculated based on Bray-Curtis distances of the CSS normalized data between samples for all time points.

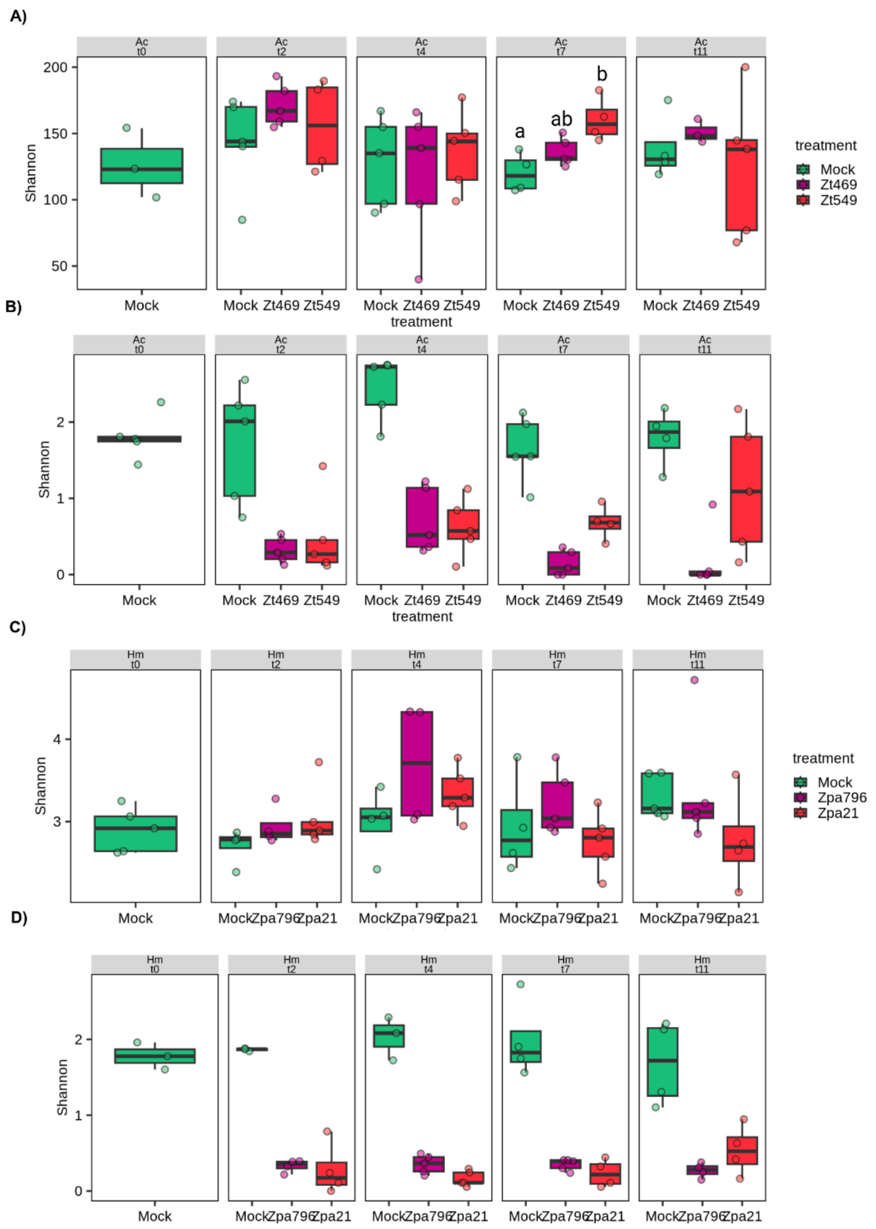


**Supplementary Figure 6. The** dynamics of the leaf microbiome of *A. cylindrica* (A, C) and *H. murinum* (B, D) infected with different lineages of *Z. tritici* and *Z. passerinii*, respectively. A, B) 16S-rRNA-V4 barcoding and C, D) ITS2 barcoding. The boxplots represent the alpha diversity (Shannon index) in the mock, virulent, and avirulent treatments by sampling time.

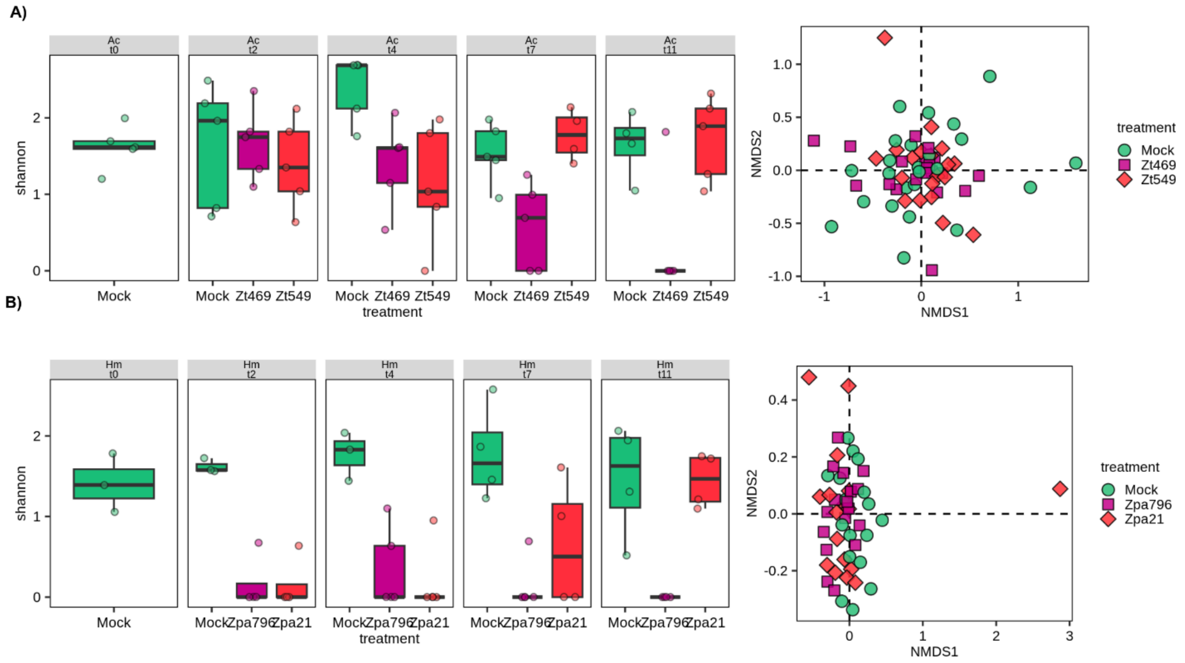


**Supplementary Figure 7.** Alpha diversity of the eukaryotic leaf microbiome of *A. cylindrica* (A) and *H. murinum* (B) infected with different lineages of *Z. tritici* and *Z. passerinii*, respectively. The *Zymoseptoria* reads were removed from the data. The boxplots represent the Shannon index by sampling point and the NMDS was defined based on Bray-Curtis distances of the CSS normalized data between treatments.

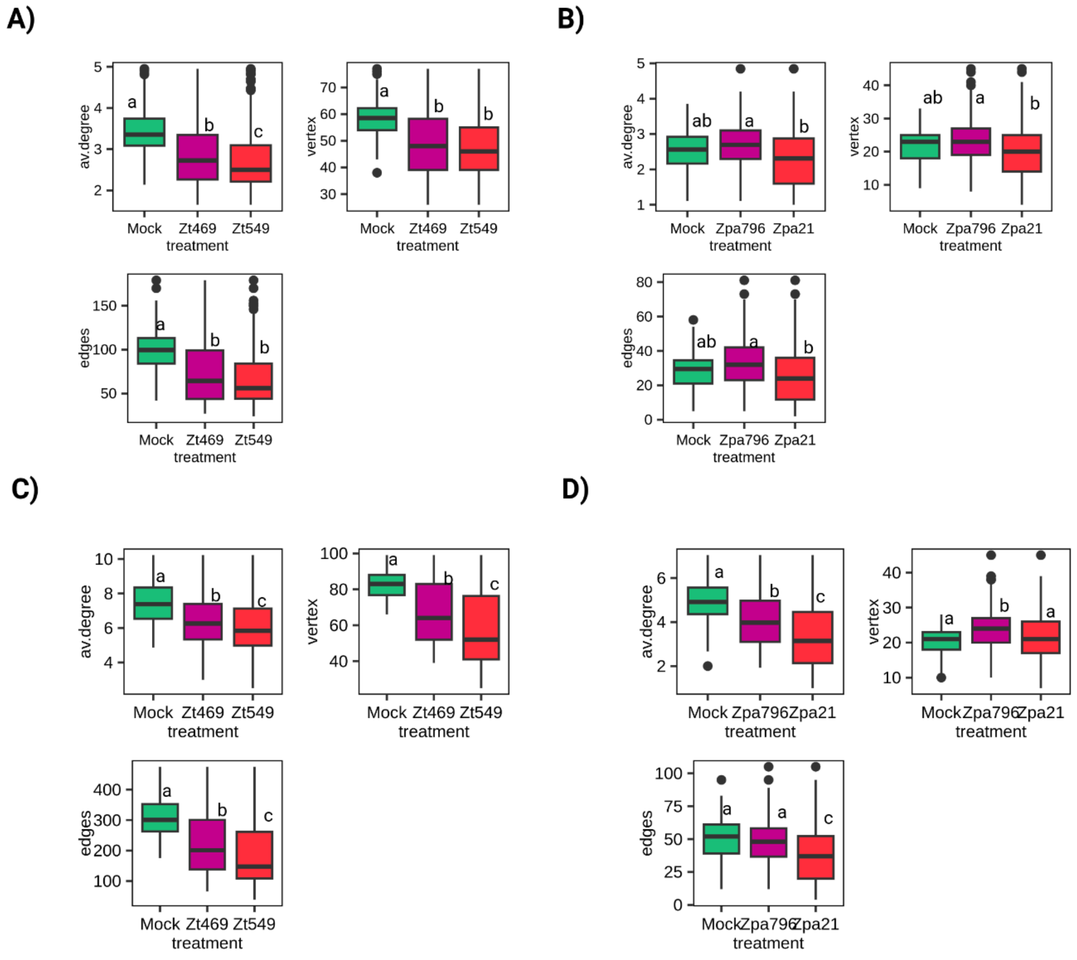


**Supplementary Figure 8.** Correlation network metrics. Average degree connectivity of OTUs (av.degree), number of correlating OTUs (vertex) and number of correlations (edges) from correlation networks of *A. cylindrica* (A, C) and *H. murinum* (B, D) using the 16S-rRNA-V4 amplicon sequence data (A, B) or the 16S-rRNA-V5V7 (C, D)

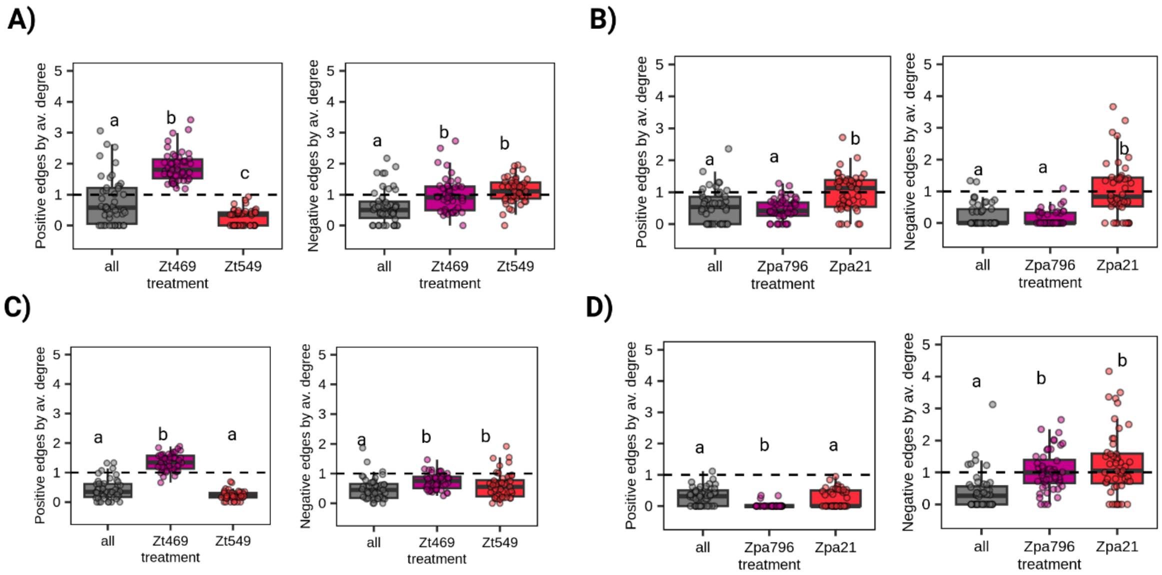


**Supplementary Figure 9.** The ratio between the number of positive and negative correlations with *Zymoseptoria* and the average degree connectivity of the OTUs from the correlation networks of *A. cylindrica* (A, C) and *H. murinum* (B, D) using the 16S-rRNA-V4 amplicon sequence data (A, B) or the 16S-rRNA-V5V7 (C, D).

A)

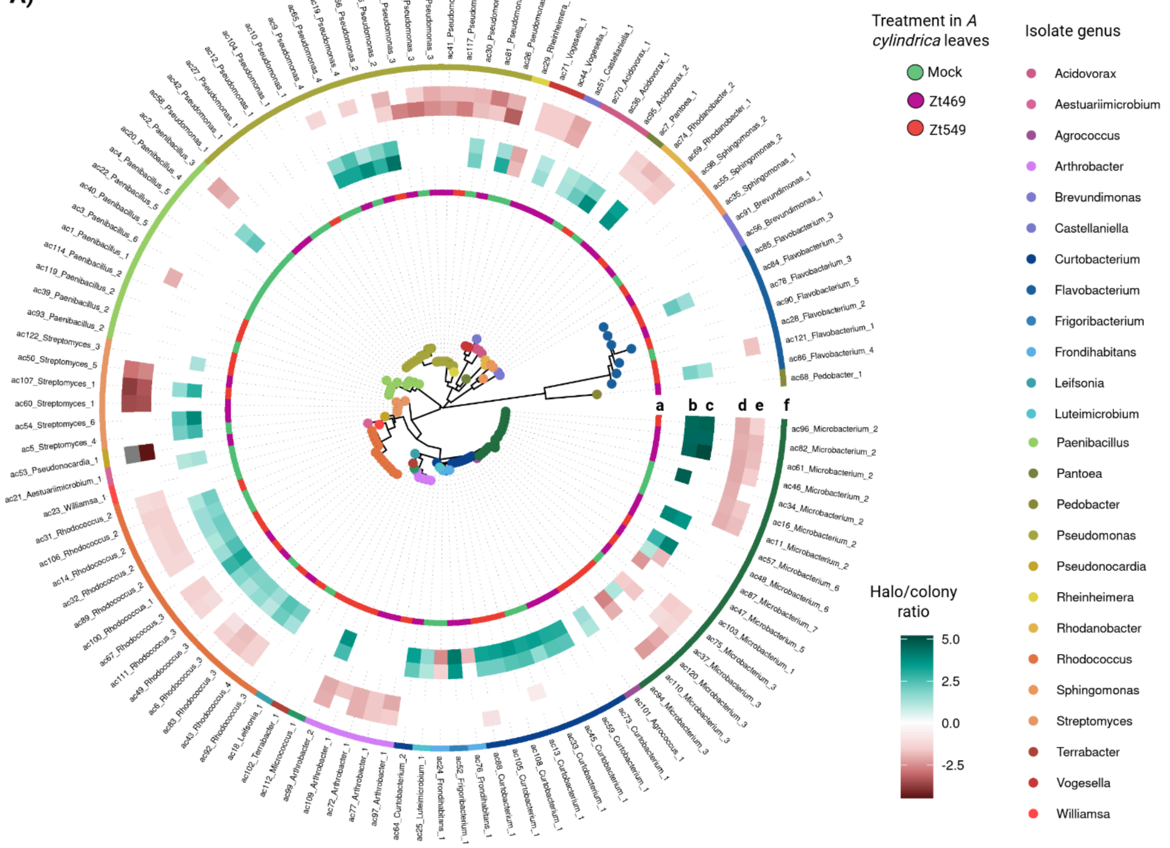

Treatment in *A cylindrica* leaves
Mock
Zt469
Zt549
Isolate genus
Acidovorax
Aestuariimicrobium
Agrococcus
Arthrobacter
Brevundimonas
Castellaniella
Curtobacterium
Flavobacterium
Frigoribacterium
Frondihabitans
Leifsonia
Luteimicrobium
Paenibacillus
Pantoea
Pedobacter
Pseudomonas
Pseudonocardia
Rheinheimera
Rhodanobacter
Rhodococcus
Sphingomonas
Streptomyces
Terrabacter
Vogesella
Williamsa
Halo/colony ratio
5.0
2.5
0.0
-2.5
a
b c
d e
f


B)

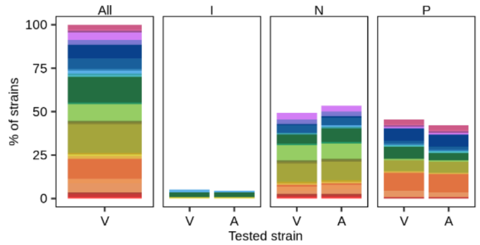

All
I
N
P
% of strains
100
75
50
25
0
V
A
Tested strain


C)

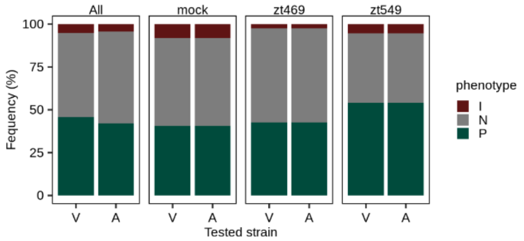

All
mock
zt469
zt549
Fequency (%)
100
75
50
25
0
V
A
Tested strain
phenotype
I
N
P


D)

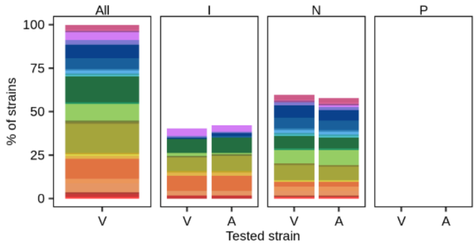

All
I
N
P
% of strains
100
75
50
25
0
V
A
Tested strain


E)

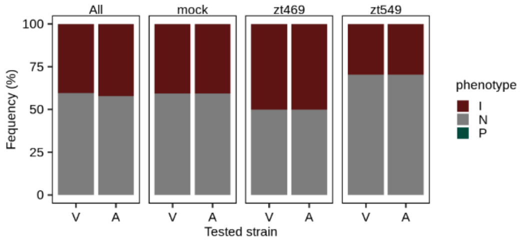

All
mock
zt469
zt549
Fequency (%)
100
75
50
25
0
V
A
Tested strain
phenotype
I
N
P

**Supplementary Figure 10.** *In-vitro* confrontation assays between the *A. cylindrica leaf-associated* bacteria and the virulent and avirulent Z. tritici Zt469 and Zt549, respectively. The heatmaps (A) show the 16s-rRNA-V4 phylogeny of the bacterial culture collection and the tracks indicate the treatment that the strains come from (a) the effect of the virulent (b) and avirulent (c) *Zymoseptoria* strain on the bacteria (B, C), the effect of the bacteria on the virulent (d) and avirulent e) *Zymoseptoria* strain (D, E) and the genus of the bacterial strain (f). The bar plots indicate the proportion of bacterial genera in the different interaction phenotypes (B, D) and the proportion across the treatment of the origin of the bacterial strain (C, E). I: growth inhibition (red). N: no effect (gray). P: growth enhancement (green)

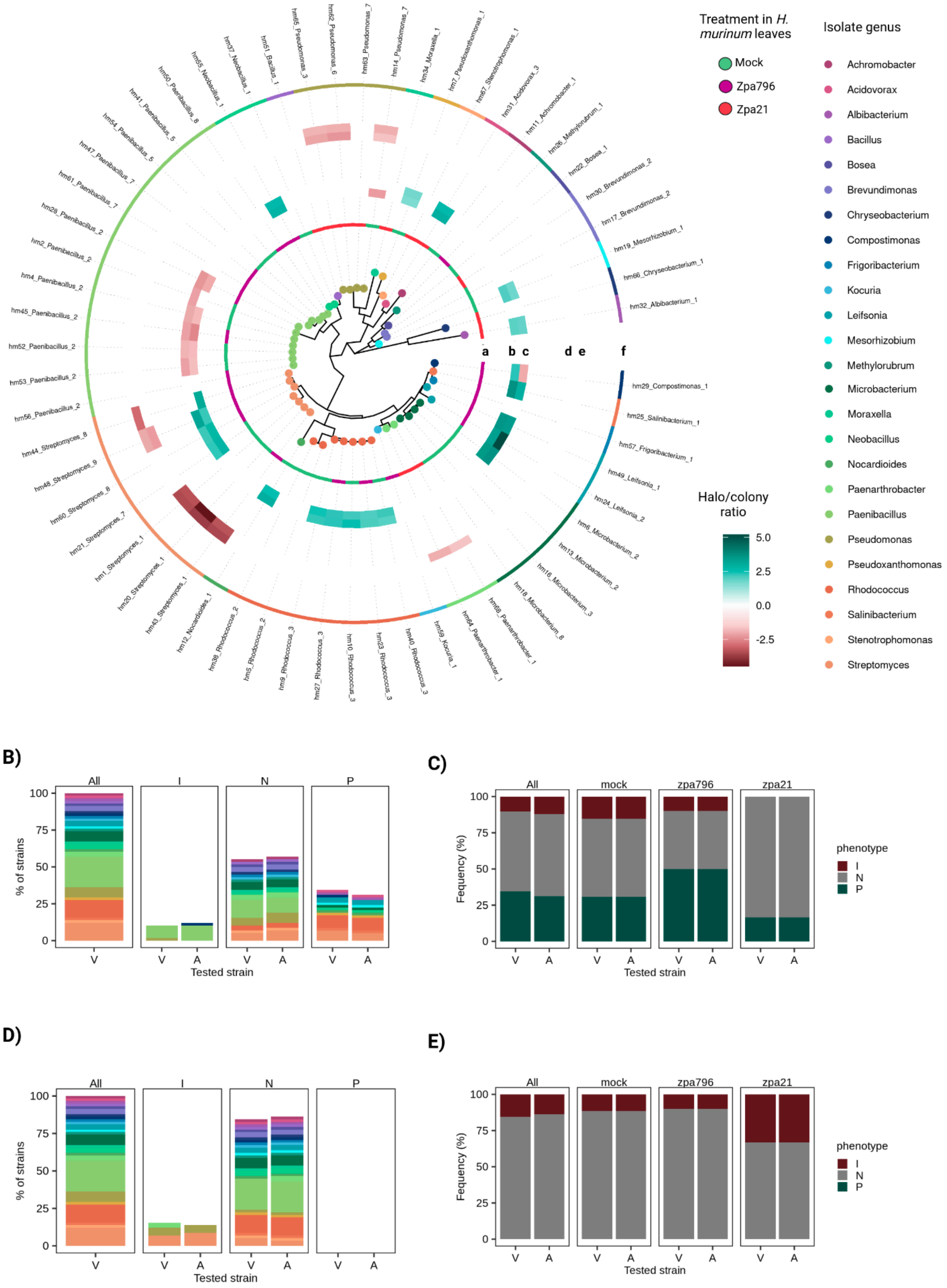

Treatment in *H. murinum* leaves
Mock
Zpa796
Zpa21
Isolate genus
Achromobacter
Acidovorax
Albibacterium
Bacillus
Bosea
Brevundimonas
Chryseobacterium
Compostimonas
Frigoribacterium
Kocuria
Leifsonia
Mesorhizobium
Methylorubrum
Microbacterium
Moraxella
Neobacillus
Nocardioides
Paenarthrobacter
Paenibacillus
Pseudomonas
Pseudoxanthomonas
Rhodococcus
Salinibacterium
Stenotrophomonas
Streptomyces
Halo/colony ratio
5.0
2.5
0.0
-2.5
a
b c
d e
f
hm65_Pseudomonas_3
hm62_Pseudomonas_6
hm63_Pseudomonas_7
hm14_Pseudomonas_7
hm34_Moraxella_1
hm7_Pseudoxanthomonas_1
hm67_Stenotrophomonas_1
hm31_Acidovorax_3
hm11_Achromobacter_1
hm26_Methylorubrum_1
hm22_Bosea_1
hm30_Brevundimonas_2
hm17_Brevundimonas_2
hm19_Mesorhizobium_1
hm66_Chryseobacterium_1
hm32_Albibacterium_1
hm29_Compostimonas_1
hm25_Salinibacterium_1
hm57_Frigoribacterium_1
hm49_Leifsonia_1
hm24_Leifsonia_2
hm6_Microbacterium_2
hm13_Microbacterium_2
hm16_Microbacterium_3
hm18_Microbacterium_8
hm68_Paenarthrobacter_1
hm64_Paenarthrobacter_1
hm59_Kocuria_1
hm40_Rhodococcus_3
hm23_Rhodococcus_3
hm10_Rhodococcus_3
hm27_Rhodococcus_3
hm9_Rhodococcus_3
hm5_Rhodococcus_2
hm36_Rhodococcus_2
hm12_Nocardioides_1
hm43_Streptomyces_1
hm20_Streptomyces_1
hm1_Streptomyces_1
hm21_Streptomyces_7
hm60_Streptomyces_8
hm48_Streptomyces_9
hm44_Streptomyces_8
hm56_Paenibacillus_2
hm53_Paenibacillus_2
hm52_Paenibacillus_2
hm45_Paenibacillus_2
hm4_Paenibacillus_2
hm2_Paenibacillus_2
hm29_Paenibacillus_2
hm61_Paenibacillus_7
hm47_Paenibacillus_7
hm54_Paenibacillus_5
hm41_Paenibacillus_5
hm50_Paenibacillus_8
hm55_Neobacillus_1
hm37_Neobacillus_1
hm51_Bacillus_1
B)
All
I
N
P
% of strains
100
75
50
25
0
V
A
Tested strain
C)
All
mock
zpa796
zpa21
Fequency (%)
100
75
50
25
0
V
A
Tested strain
phenotype
I
N
P
D)
All
I
N
P
% of strains
100
75
50
25
0
V
A
Tested strain
E)
All
mock
zpa796
zpa21
Fequency (%)
100
75
50
25
0
V
A
Tested strain
phenotype
I
N
P

**Supplementary Figure 11.** *In-vitro* confrontation assays between the *H. murinum leaf-associated* bacteria and the virulent and avirulent Z. passerinii Zpa796 and Zpa21, respectively. The heatmaps (A) show the 16s-rRNA-V4 phylogeny of the bacterial culture collection and the tracks indicate the treatment that the strains come from (a) the effect of the virulent (b) and avirulent (c) *Zymoseptoria* strain on the bacteria (B, C), the effect of the bacteria on the virulent (d) and avirulent e) *Zymoseptoria* strain (D, E) and the genus of the bacterial strain (f). The bar plots indicate the proportion of bacterial genera in the different interaction phenotypes (B, D) and the proportion across the treatment of the origin of the bacterial strain (C, E). I: growth inhibition (red). N: no effect (gray). P: growth enhancement (green).

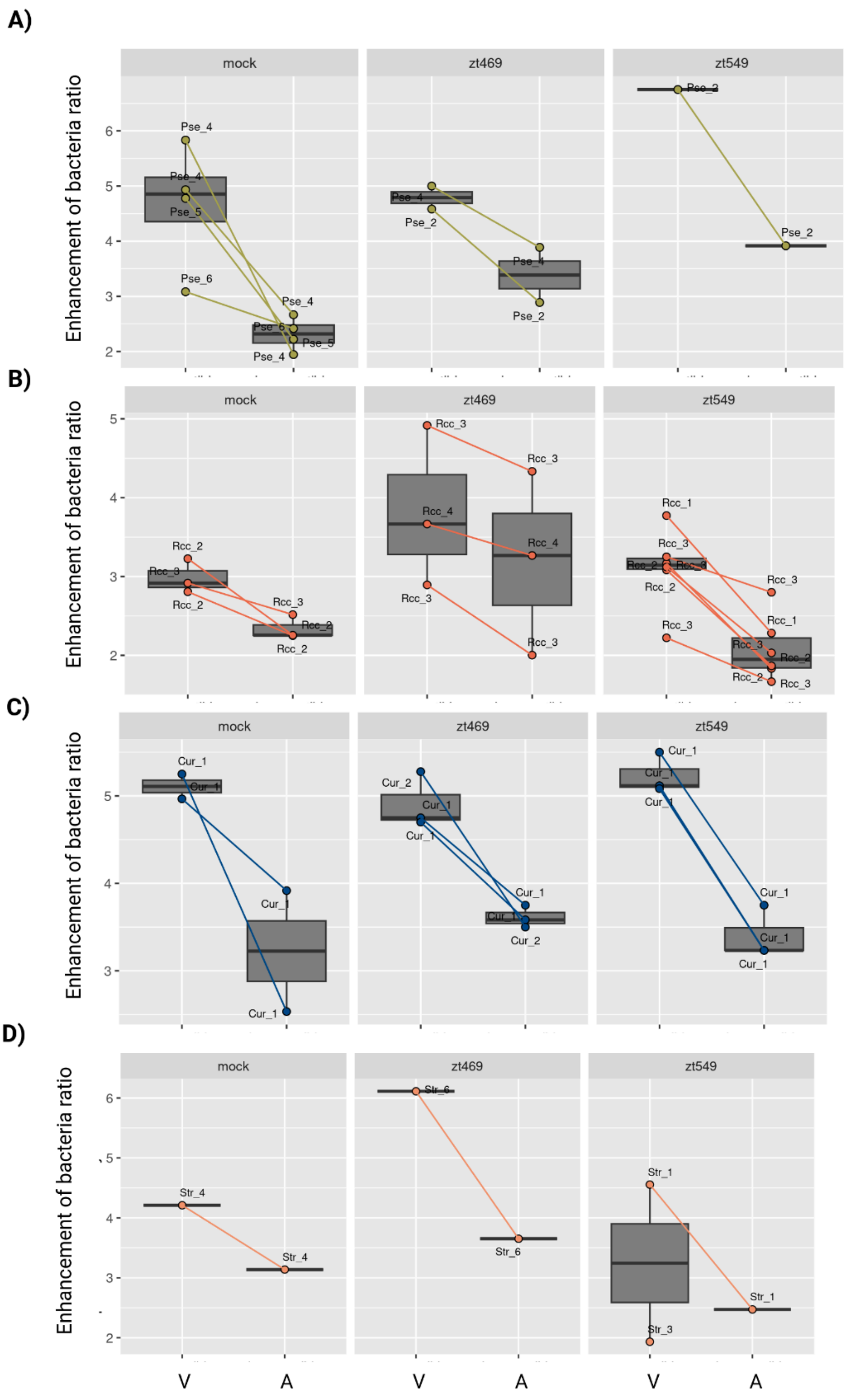


**Supplementary Figure 12.** Growth enhancement phenotype induced by virulent (V) and avirulent (A) *Z. tritici* Zt469 and Zt549 in different bacterial phylotypes. The strains are separated by the treatment of origin. A) *Pseudomonas* phylotypes, B) *Rhodococcus phylotypes*, C) *Curtobacterium* phylotypes, and D) *Streptomyces* phylotypes.

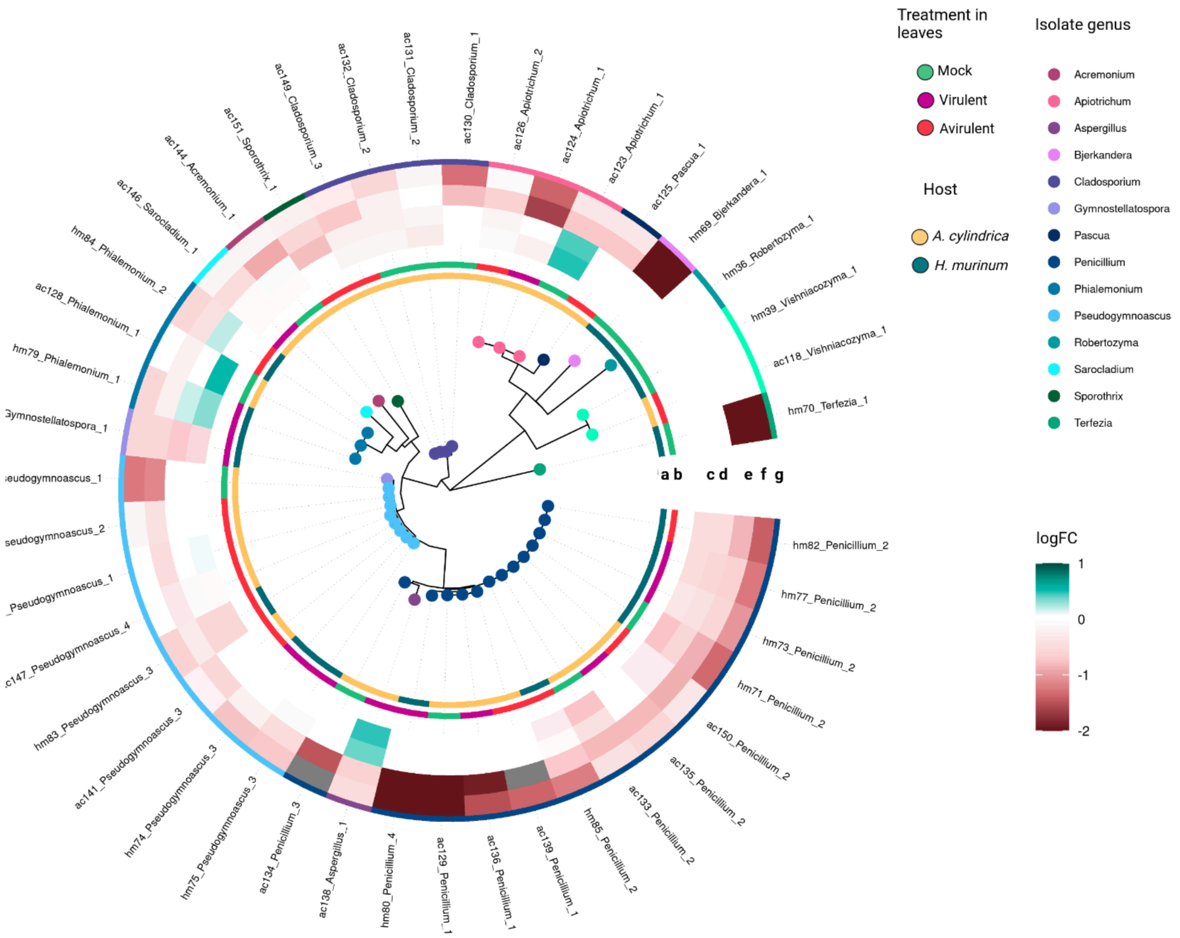


**Supplementary Figure 13.** *In-vitro* confrontation assays between the leaf-associated fungi of A. cylindrica and *H. murinum* and the virulent and avirulent Z. tritici (Zt469 and Zt549 respectively) and Z. passerinii (Zpa796 and Zpa21, respectively). The heatmaps show the ITS2 phylogeny of the fungi culture collection and the tracks indicate the host (a) and treatment (b) that the strains come from, the effect of the virulent (c) and avirulent (d) *Zymoseptoria* strain on the fungi, the effect of the fungi on the virulent (e) and avirulent (f) *Zymoseptoria* strain and the genus of the bacterial strain (g). I: growth inhibition (red). N: no effect (gray). P: growth enhancement (green).

### References


1. Fagundes WC, Haueisen J, Stukenbrock EH. Dissecting the Biology of the Fungal Wheat Pathogen *Zymoseptoria tritici* : A Laboratory Workflow. Curr Protoc Microbiol [Internet]. 2020 Dec [cited 2022 Jul 19];59(1). Available from: https://onlinelibrary.wiley.com/doi/10.1002/cpmc.128

2. Rojas-Barrera IC, Flores-Núñez VM, Haueisen J, Alizadeh A, Salimi F, Stukenbrock EH. Evolution of sympatric host-specialized lineages of the fungal plant pathogen Zymoseptoria passerinii in natural ecosystems. New Phytol. 2025;245(4):1673–87.

3. Seybold H, Demetrowitsch TJ, Hassani MA, Szymczak S, Reim E, Haueisen J, et al. A fungal pathogen induces systemic susceptibility and systemic shifts in wheat metabolome and microbiome composition. Nat Commun. 2020 Dec;11(1):1910.

4. Gohl D, Gohl DM, MacLean A, Hauge A, Becker A, Walek D, et al. An optimized protocol for high-throughput amplicon-based microbiome profiling. Protoc Exch [Internet]. 2016 Jul 25 [cited 2025 May 28]; Available from: https://www.protocols.io/view/an-optimized-protocol-for-high-throughput-amplicon-n92ldr4oxg5b/v1

5. Flores-Núñez VM, Camarena-Pozos DA, Chávez-González JD, Alcalde-Vázquez R, Vázquez-Sánchez MN, Hernández-Melgar AG, et al. Synthetic communities increase microbial diversity and productivity of Agave tequilana plants in the field. Phytobiomes J. 2023;7(4):435–48.

6. Bolger AM, Lohse M, Usadel B. Trimmomatic: a flexible trimmer for Illumina sequence data. Bioinformatics. 2014;30(15):2114–20.

7. Rognes T, Flouri T, Nichols B, Quince C, Mahé F. VSEARCH: a versatile open source tool for metagenomics. PeerJ. 2016;4:e2584.

8. Wang Q, Cole JR. Updated RDP taxonomy and RDP Classifier for more accurate taxonomic classification. Microbiol Resour Announc. 2024;13(4):e01063-23.

9. Abarenkov K, Zirk A, Piirmann T, Pöhönen R, Ivanov F, Nilsson RH, et al. UNITE USEARCH/UTAX release for eukaryotes [Internet]. UNITE Community; 2022 [cited 2025 May 28]. Available from: https://doi.plutof.ut.ee/doi/10.15156/BIO/2483924

10. R Core Team. R: A Language and Environment for Statistical Computing [Internet]. Vienna, Austria: R Foundation for Statistical Computing; 2021. Available from: https://www.R-project.org/

11. Wickham H. ggplot2: Elegant Graphics for Data Analysis [Internet]. Springer-Verlag New York; 2016. Available from: https://ggplot2.tidyverse.org

12. Oksanen J, Simpson GL, Blanchet FG, Kindt R, Legendre P, Minchin PR, et al. vegan: Community Ecology Package [Internet]. 2024. Available from: https://CRAN.R-project.org/package=vegan

13. Paulson JN, Stine OC, Bravo HC, Pop M. Differential abundance analysis for microbial marker-gene surveys. Nat Methods. 2013 Dec;10(12):1200–2.

14. Paulson JN, Pop M, Bravo HC. metagenomeSeq: Statistical analysis for sparse high-throughput sequencing. Bioconductor Package. 2013;1(0):191.

15. McMurdie PJ, Holmes S. phyloseq: An R package for reproducible interactive analysis and graphics of microbiome census data. PLoS ONE. 2013;8(4):e61217.

16. Lahti L, Shetty S. microbiome R package. 2012.

17. Gao CH, Dusa A. ggVennDiagram: A “ggplot2” Implement of Venn Diagram [Internet]. 2024. Available from: https://CRAN.R-project.org/package=ggVennDiagram

18. Gehlenborg N. UpSetR: A More Scalable Alternative to Venn and Euler Diagrams for Visualizing Intersecting Sets [Internet]. 2019. Available from: https://CRAN.R-project.org/package=UpSetR

19. Friedman J, Alm EJ. Inferring Correlation Networks from Genomic Survey Data. Von Mering C, editor. PLoS Comput Biol. 2012 Sep 20;8(9):e1002687.

20. Kurtz Z, Mueller C, Miraldi E, Bonneau R. SpiecEasi: Sparse Inverse Covariance for Ecological Statistical Inference [Internet]. 2024. Available from: https://github.com/zdk123/SpiecEasi

21. Csardi G, Nepusz T. The igraph software. Complex Syst. 2006;1695:1–9.

22. Schloerke B, Cook D, Larmarange J, Briatte F, Marbach M, Thoen E, et al. GGally: Extension to “ggplot2” [Internet]. 2024. Available from: https://CRAN.R-project.org/package=GGally

23. Lin H, Peddada SD. Analysis of microbial compositions: a review of normalization and differential abundance analysis. Npj Biofilms Microbiomes. 2020 Dec 2;6(1):60.

24. Lyda TA, Joshi MB, Andersen JF, Kelada AY, Owings JP, Bates PA, et al. A unique, highly conserved secretory invertase is differentially expressed by promastigote developmental forms of all species of the human pathogen, Leishmania. Mol Cell Biochem. 2015;404:53–77.

25. Emms, D. M., & Kelly, S. (2015). OrthoFinder: solving fundamental biases in whole genome comparisons dramatically improves orthogroup inference accuracy. Genome Biology, 16(157), 1–14. https://doi.org/10.1186/s13059-015-0721-2

26. Buchfink, B., Xie, C., & Huson, D. H. (2014). Fast and sensitive protein alignment using DIAMOND. Nature Methods, 12(1), 59–60. https://doi.org/10.1038/nmeth.3176

27. Lapalu, N., Lamothe, L., Petit, Y., Genissel, A., Delude, C., Feurtey, A., Abraham, L. N., Smith, D., King, R., Renwick, A., Appertet, M., Sucher, J., Steindorff, A. S., Goodwin, S. B., Kema, G. H. J., Grigoriev, I. V., Hane, J., Rudd, J., Stukenbrock, E., … Lebrun, M.-H. (2023). Improved gene annotation of the fungal wheat pathogen Zymoseptoria tritici based on combined Iso-Seq and RNA-Seq evidence. https://doi.org/10.1101/2023.04.26.537486

28. Feurtey, A., Lorrain, C., Croll, D., Eschenbrenner, C., Freitag, M., Habig, M., Haueisen, J., Möller, M., Schotanus, K., & Stukenbrock, E. H. (2020a). Genome compartmentalization predates species divergence in the plant pathogen genus Zymoseptoria. BMC Genomics, 21(1). https://doi.org/10.1186/s12864-020-06871-w

29. Wyatt, N. A., Spanner, R. E., & Bolton, M. D. (2024). The Complete and Gapless Genome Sequence of the Sugarbeet Pathogen Cercospora beticola . PhytoFrontiersTM, 4(3), 434–437. https://doi.org/10.1094/phytofr-11-23-0146-a

30. Martin, M. (2011). Cutadapt removes adapter sequences from high-throughput sequencing reads. EMBnet.Journal, 17(1), 10–12. https://doi.org/https://doi.org/10.14806/ej.17.1.200

31. Bankevich, A., Nurk, S., Antipov, D., Gurevich, A. A., Dvorkin, M., Kulikov, A. S., Lesin, V. M., Nikolenko, S. I., Pham, S., Prjibelski, A. D., Pyshkin, A. V., Sirotkin, A. V., Vyahhi, N., Tesler, G., Alekseyev, M. A., & Pevzner, P. A. (2012). SPAdes: A new genome assembly algorithm and its applications to single-cell sequencing. Journal of Computational Biology, 19(5), 455–477. https://doi.org/10.1089/cmb.2012.0021

32. Mikheenko, A., Prjibelski, A., Saveliev, V., Antipov, D., & Gurevich, A. (2018). Versatile genome assembly evaluation with QUAST-LG. Bioinformatics, 34(13), i142–i150. https://doi.org/10.1093/bioinformatics/bty266

33. Simão, F. A., Waterhouse, R. M., Ioannidis, P., Kriventseva, E. V., & Zdobnov, E. M. (2015). BUSCO: Assessing genome assembly and annotation completeness with single-copy orthologs. Bioinformatics, 31(19), 3210–3212. https://doi.org/10.1093/bioinformatics/btv351

34. Brůna, T., Hoff, K. J., Lomsadze, A., Stanke, M., & Borodovsky, M. (2021). BRAKER2: Automatic eukaryotic genome annotation with GeneMark-EP+ and AUGUSTUS supported by a protein database. NAR Genomics and Bioinformatics, 3(1), 1–11. https://doi.org/ 10.1093/nargab/lqaa108

35. Ware SB, Verstappen ECP, Breeden J, Cavaletto JR, Goodwin SB, Waalwijk C, et al. Discovery of a functional Mycosphaerella teleomorph in the presumed asexual barley pathogen Septoria passerinii. Fungal Genet Biol. 2007 May;44(5):389–97
